# Tool Choice drastically Impacts CRISPR Spacer-Protospacer Detection

**DOI:** 10.1101/2025.05.06.652306

**Authors:** Uri Neri, Antonio Pedro Camargo, Brian Bushnell, Simon Roux

## Abstract

CRISPR (Clustered Regularly Interspaced Short Palindromic Repeats) systems are a fundamental defense mechanism in prokaryotes, where short sequences called spacers are stored in the host genome to recognize and target exogenous genetic elements. Viromics, the study of viral communities in environmental samples, relies heavily on identifying these spacer-target interactions to understand host-virus relationships. However, the choice of sequence search tool to identify putative spacer targets is often overlooked, leading to an unknown impact of downstream inferences in virus-host analysis. Here, we utilize simulated and real datasets to compare popular sequence alignment and search tools, revealing critical differences in their ability to detect multiple matches and handle varying degrees of sequence identity between spacers and potential targets. Finally, we provide general guidelines that may inform future research regarding matching, which is a common practice in studying the complex nature of host-MGE interactions.

## 2 Introduction

CRISPR (clustered regularly interspaced short palindromic repeats) systems play a vital role in prokaryotic defense against mobile genetic elements, including viruses, plasmids, and other autonomous genetic elements [19, 11]. These systems are organized as arrays in the bacteria or archaea genome, where short sequences called spacers are interspersed between repeated sequences. The spacer sequences within these arrays guide the targeting of invasive genetic elements, allowing for specific defense against these threats [16]. The corresponding locus on the virus genome where the spacer complements is termed “protospacer”. The analysis of spacer-protospacer pairs is essential in understanding the complex interactions between hosts and MGEs [10].

The identification of genuine host-MGE interactions through spacer-protospacer matching presents unique challenges due to the dynamic nature of these relationships and the complexity of sequence evolution. While matches between spacers and protospacers are often interpreted as evidence of interaction, various biological and technical factors can complicate this interpretation [10, 29].

Several key scenarios can lead to false positive assignments in spacer-protospacer matching. Low complexity sequences can create spurious matches between simple repeat regions (albeit these can be mitigated through complexity filtering via Shannon entropy or DUST). Another type of potential false positives are highly conserved sequences shared by unrelated MGEs, potentially resulting from horizontal gene transfer between MGEs. The horizontal transfer of CRISPR arrays themselves on mobile elements further requires careful examination of array genomic context (regions outside the CRISPR loci) and phylogenetic analysis. Self-targeting events, where matches occur against the host genome rather than MGEs, necessitate comparison against host genome databases and analysis of targeting context. Finally, historical acquisition events may not reflect current interactions, requiring consideration of phylogenetic dating, evolution rates and the effects of the protospacers being under selective pressure to mutate (which may reduce the MGE susceptibility to deterioration by the CRISPR system). This is further complicated by the fact that increasing the allowed distance between sequences directly increases the likelihood of considering sequences similar or related to each other.

False negatives present another challenge in spacer-protospacer matching, particularly when dealing with large databases of potential targets. Many alignment and search tools default to reporting only the best matches or the first matches that pass a given threshold for a given query or HSP. This may result in potentially missing additional legitimate matches. Unfortunately, different tools also handle ambiguous or secondary alignments differently: they may be reported, reported up to a number or based on relative alignment quality, or omitted. Similarly, cases where a query sequence has multiple matches in different reference sequences are not handled uniformly across tools. This limitation becomes increasingly problematic as databases grow larger and more diverse, a single spacer might match (implying a targeting) multiple related MGEs.

Yet despite these variations, the choice of spacer-to-protospacer search or alignment tool is often not deeply considered. Presently, the common option for this task, popularized by Edwards et al and Biswas et al [10, 2], uses BLASTn [1] with parameters adjusted for short input sequences. However for most bioinformatic tools, the exact workflow design and parameter choice can impact the outcome, including in sequence analysis. The importance of proper tool usage and parameter interpretation is highlighted by historical examples in bioinformatics. A striking example is the work of Shah et al, Shah et al. [26], in which they report how certain misunderstandings of BLAST’s `-max_target_seqs` parameter may lead to incorrect assumptions about result completeness, potentially impacting published analyses. Albeit this was later clarified by Madden et al., [14] (of the blast development team) as an unfortunate combination of a software bug (that were since patched) affecting rare cases, and misconceptions regarding the process BLAST+ uses for tie-breaking (alignments of equal plausibility), and finally a consideration regarding composition base scoring. Apart from the patched bug, the main outcome of this correspondence led to more explicit details in blast documentation (specifically the appendix “Outline of the BLAST process”). Still, this highlights that misconceptions about the expected exhaustiveness of tools’ result-reporting can also lead to incorrect assumptions about the outcome of an analysis. In practice, most bioinformatic tools use various heuristics and optimizations, typically designed with specific use cases in mind. For example, most short-read mappers assume the reference to be the output of a singular assembly - which would imply the reference does not contain extremely redundant copies of the same loci, or a limited number of very similar sequences (e.g. strain variants, alternative splice variants), and this assumption impacts the way read mapping is computed and results are reported.

The choice of tool and its parameters can significantly impact the detection of these multiple matches, with some tools prioritizing speed over completeness by limiting the number of reported matches, or by other internal heuristics such as seed sequence selection from high occurring sequences being penalized. This trade-off between sensitivity and computational efficiency is especially important to consider as most available tools were designed for different tasks than spacer-protospacer matching (e.g. expression analysis, homology detection, and variant calling), and under different assumptions (such as reference and query sequence size and database size or the nature of the reference source: from a single isolate or metagenomic sample rather than from aggregation of sequences from different sources).

Another potential consideration is the computational resources requirements. Memory, storage, and availability of CPU cores are factors differing between tools. Parameter choice may also impact these factors considerably, with certain tools offering tunable parameters to trade-off between sensitivity and computational efficiency. In recent years, spacer database size has been rapidly increasing - from 366,799 unique spacers in 2017 [28] to 1,173,006 unique spacers reported in 2021 [8] to 3,835,942 unique spacers in 2023 [5]. Similarly, public virus and MGE databases are growing rapidly, with large contributions from metagenomic samples resulting in routine fold increases in the number of predicted viral contigs [5]. Most tools require more resources as the size of the database grows, and as this trend continues, certain workflows and tools may become prohibitively expensive to run in a reasonable time frame.

## 3 Methods

### 3.1 Tool Selection

We evaluated several widely-used sequence alignment and search tools, alongside an implementation of basic string containment. The tools were selected based on their availability and historical use, and were chosen to span a range of different algorithmic approaches. It is important to note that these tools were not specifically designed for spacer-protospacer matching, but rather for more general sequence search tasks (mmseqs2, blastn-short and lexicmap), or alignment/mapping of short reads to reference genomes (bowtie1, bowtie2, minimap2, nucmer, bbmap-skimmer, strobealign). Additionally, our focus here is specifically on spacer-protospacer matching, and so we did not evaluate general host-prediction tools (even those that may perform spacer-protospacer matching), like SpacePHARER [35] or iPHoP [24], which may perform additional analyses based on prior additional information, such as evaluating the LCA from multiple spacer-protospacer matches.

### 3.2 Data Generation and acquisition

### 3.3 Synthetic dataset

In order to examine each tool’s performance on diverse spacer-to-target matches scenario (i.e. different number of match, similarity between spacer and target, etc), we created a simplistic command-line interface Python script automating the simulation of spacer-protospacer pairs given a set of parameters (see code and data availability). These parameters include target contig length, GC content, spacer length, number of occurrences in the target(s), range of mismatches, reverse complement frequency. This allows for fine grain control over the simulated spacer and target sequences, while recording the ground truth for later validation. See supplementary figure 1 for a visual representation of the data generation workflow.

### 3.4 Real datasets

To evaluate the same tools in a “real-life” scenario, We used the predicted viral contigs and CRISPR spacers from IMG/VR4 [5]. IMG/VR4 was chosen for several reasons: 1) it is one of the most comprehensive databases of uncultured phage and viral genomes, and 2) the associated CRISPR database contains a large number of CRISPR spacer data from various genomes and metagenomes, 3) the original database released included an existing attempt at spacer-protospacer matching using one of the tools evaluated (blastn, albeit with different parameters, namely the “–max-target-seqs”), allowing for a direct comparison to historical results. Briefly, the viral contigs in IMG/VR4 are predicted using a combination of tools (primarily genomad [4]) supplemented with existing databases such as NCBI’s RefSeq and GenBank, while the CRISPR spacers were compiled from previously predicted CRISPR arrays identified in IMG/M genomes and metagenomes (primarily via piler-cr [9] and CRT [3]). For more information, please refer to the IMG/VR4 paper [5].

### 3.5 Alignment and Recalculation of Mismatches

For comparing alignments across tools, we used the Levenshtein distance [13] (a.k.a edit distance) as our main scoring metric. For any two sequences, this metric sums both substitutions and gaps (i.e. insertions and deletions) into a single value. This choice was made to ensure consistent comparison across tools that may handle gaps differently, and because in the context of CRISPR-based defense, both substitutions and gaps can affect the efficiency of target recognition.

To ensure consistent mismatch calculation across tools, we realigned all reported matches using parasail [7], specifically the `nw_stats_scan` i.e. Needleman-Wunsch global alignment with gap opening and extension penalties of 10, and a scoring matrix of `nuc44`. This allowed us to:

1. Calculate edit distances independently of tool-specific scoring schemes
2. Verify reported matches and their orientations
3. Generate standardized alignment visualizations for manual comparison

### 3.6 Performance definition and calculation

We defined true and false positives/negatives differently for synthetic and real datasets. For synthetic datasets, sequences were generated following pre-planned insertion patterns with known coordinates, strands, and number of mismatches. While analyzing 7,638,511 pre-planned spacer occurrences, we discovered that some sequences could align at unplanned locations while still meeting our mismatch threshold (3). To ensure accuracy, we recalculated all alignments and filtered out sequences with extracted contig regions longer than 130bp (120bp being the maximum spacer length). After this validation, we identified 999,916 additional valid alignments (13 with 0 mismatches, 68,612 with 1 mismatch, 224,044 with 2 mismatches, and 707,247 with 3 mismatches), demonstrating that with short sequences, the number of potential spurious alignments increases substantially with the allowed mismatch threshold. Given their validated alignment scores, we included both pre-planned and these additional alignments in our positive set. For real data (IMG VR4), the positive set comprised all alignments from all tools that met the specified threshold criteria. In both cases, we calculated standard performance metric:

Recall = true positives / (true positives + false negatives)

where true positives are matches found by a tool that exist in the positive set, false positives are matches reported by a tool that do not exist in the positive set, and false negatives are matches in the positive set that were not found by the tool. We note that in this context, “true negative” (i.e., a false alignment a tool did not report) does not represent a meaningful metric.

### 3.7 Benchmarking framework

### 3.8 Computational Resource and runtime tracking

While we have made the runtime and resource (memory and storage usage) data available, the primary focus of this study was to evaluate the ability of each tool to accurately identify spacer-protospacer matches, rather than their computational performance. However, as resource usage may become a limiting factor in the future (with the advent and accumulation of metagenomic sequence data), we designed the benchmark to automatically log several usage metrics.

For the synthetic data, we used hyperfine [23] to track runtime and resource usage, with a maximum of 5 runs for each tool. For the real data, we used SLURM’s built-in accounting system for cluster-based runs (`sacct`) via a custom script (pyseff.py)[22]. For each tool, several metrics were recorded, including wall clock time, Peak memory usage, CPU utilization, I/O operations (see fig-workflow for an overview of the synthetic data workflow). Memory efficiency was calculated as the ratio of the peak memory usage divided by the total allocated memory, and it is important to note that certain tools would identify available memory and utilize all of it, while other tools would utilize it in a deterministic manner given input size, parameters, threads, batch size, etc. This can also affect the overall runtime for different tools, making it difficult to directly compare the scaling efficiency of different tools. Additionally, the runtime reported is not the actual runtime of the tool, but rather the runtime of the SLURM job. While most tools were run as a single job, some tools (notably blastn-short) failed to complete given our maximum available resources (either due to time or memory constraints), and were instead run as multiple jobs, on different subsets of the data. As such, the runtime reported for these tools is the sum of the runtime of all their jobs.

### 3.9 Versioning and Reproducibility

All tools were installed and managed using conda [6] (via the mamba [17] package manager) in isolated environments. Each tool was installed in a separate environment to prevent dependency conflicts and ensure reproducibility. Environment activation time was excluded from performance measurements to focus on actual tool runtime.

The exact versions and configurations of all tools were recorded in environment files, allowing for exact replication of our testing environment. All benchmarks were performed on identical hardware configurations to ensure fair comparison.

### 3.10 Extensibility

The framework is designed to be expandable through the integration of new tools. Each tool/software configuration is saved as a separate JSON file, which includes the exact commands and conda/mamba environment it uses. This configuration files can use placeholder variables which the main benchmarking script replaces with user choices during execution (such as {threads}, {contigs_file}, {spacers_file}, {output_dir}, and {results_dir}). A new JSON file can be added manually or via bench.utils.tool_commands:add_tool function in a semi automated method.

## 4 Results

### 4.1 Performance as a function of mismatch threshold

First we investigated potential tradeoffs and effects of the total edit distance (henceforth, interchangeable with mismatches) on the observed recall metric of the tools. Generally, the detection rate of each tool decreases as mismatch thresholds increase. Additionally, no single tool was able to identify all spacer occurrences, although at 0 mismatches the recall of bowtie1, bowtie2, blastn and mummer4 is approximately 0.99 (See supplementary table 2 for the recall values for each tool at different mismatch thresholds). At increased allowed mismatches, the tools showed more divergence, yet bowtie1 remained the single tool with the most unique matches by a considerable margin (Figure 1). Overall, the performance of the tools is similar between the synthetic and real datasets, albeit the overall lower sample size of the synthetic data should be considered when interpreting the results (see table 1).

**Figure 1:**
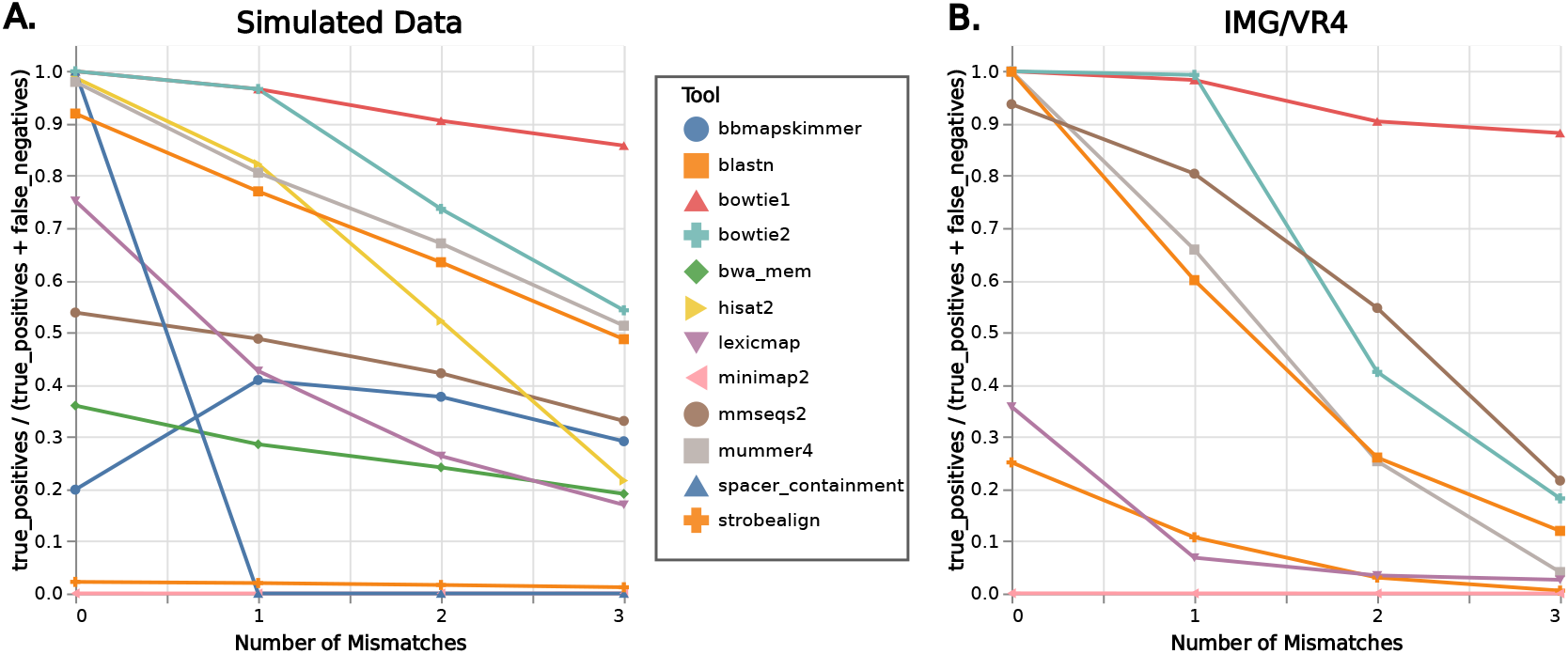
Performance of each tool as a function of mismatch threshold. The horizontal axis shows the number of allowed mismatches, while the vertical axis represents the mean detection fraction (0-1) aggregated across all spacer-contig pairs at a given mismatch threshold. Each color and shape indicates a different tool plot (shapes connected by lines for interpolation) Panel B shows the performance of the tools on the IMG/VR4 dataset, while panel A shows the performance of the tools on the synthetic dataset.

**Table 1:** Evaluated Tools and Their Characteristics. *Indexing:* “Optional” - *A persistent index can be generated (and used across multiple mapping/alignment runs) but a one-off command can also do this ‘on the fly’*. “Yes”* - A precomputed index is required, *“No”* - Indexing or building the database is not required. All tools were run with default parameters unless otherwise specified. As a general rule, parameters for each tool were chosen to maximize sensitivity or avoid limiting the ability to detect multiple matches. For more details about the exact configuration and version of each tool, see the code availability below or supplementary table 1.

| Aligner | Indexing* | Publication Year | Notes |
| --- | --- | --- | --- |
| Bowtie1 | Yes | 2009 | Last updated 2017, optimized for short sequences (25-50 bp, max 1kbp) |
| Bowtie2 | Yes | 2012 | FM Index, supports longer reads (>50bp) than v1. |
| Minimap2 | Optional | 2018 | Versatile, supports various sequence types |
| BBMap-skimmer | Optional | 2014 | Part of BBTools suite |
| StrobeAlign | Yes | 2022 | strobemer-based indexing. |
| BLAST+ | Optional | 2009 | BLASTN-short is a mode (task parameter), optimized for short sequences. |
| MMseqs2 | Optional | 2017 | Fast protein/nucleotide search |
| spacer-containment | No | - | Simple string containment. FASTA parsing with needletail, parallel processing with rayon, and string matching with memchr::memmem::Finder |
| mummer4 | Optional | 2005 | Fast protein/nucleotide search |
| LexicMap | Yes | 2024 | **Preprint** - Highly scalable, designed for fast querying of genes, genomes and long reads (>100bp) |

**Table 2:**
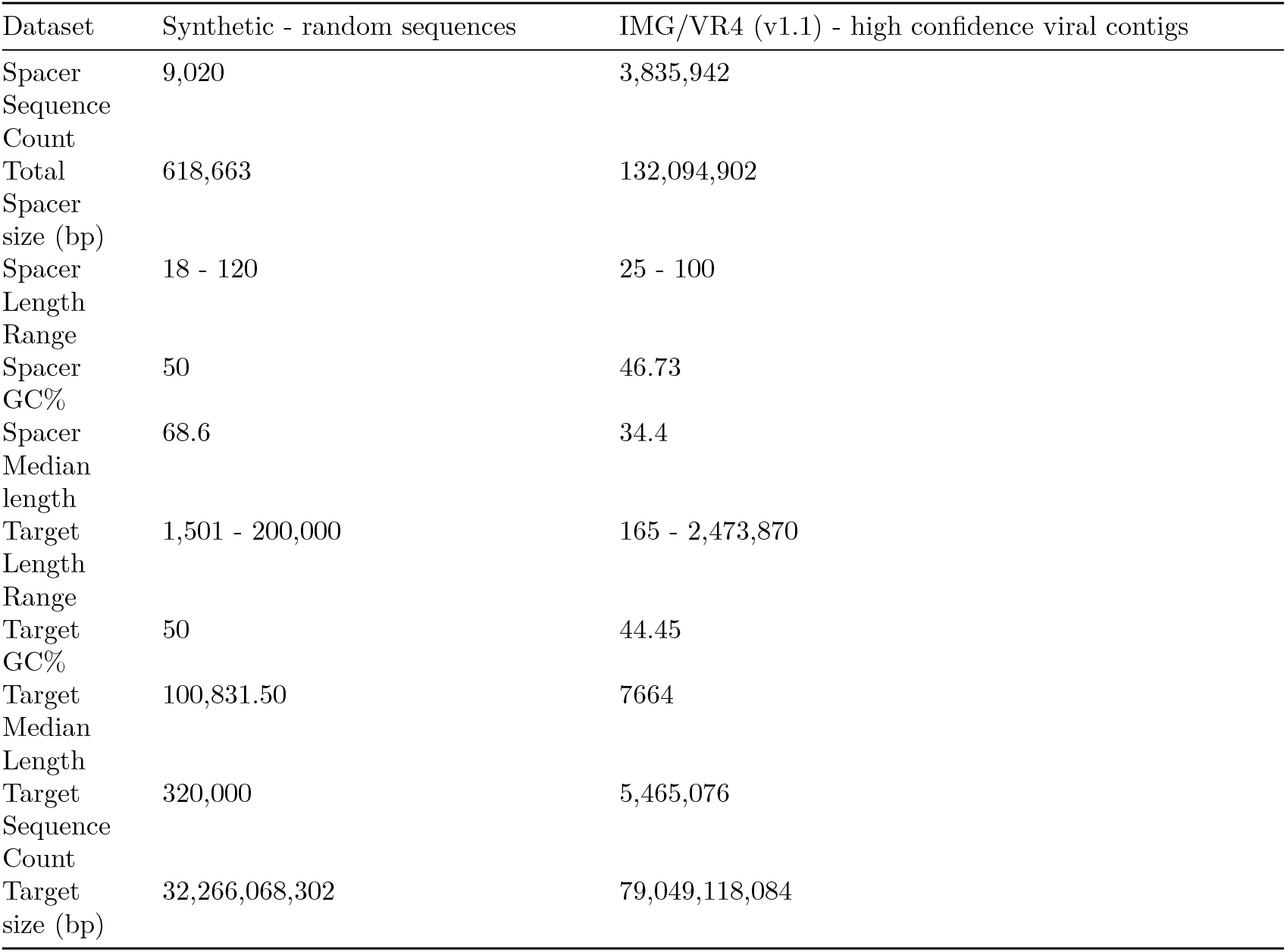
Dataset characteristics. Target is interchangeable with “contig”.

### 4.2 Performance as a function of query (spacer) abundance in reference database

We then investigated if there are any potential effects for the number of times each protospacer sequence appears in the target set (i.e. the virus sequence set). For perfect matches (0 mismatches), bowtie1 demonstrates exceptional performance with recall rates consistently above 0.99 across all occurrence frequencies (Figure 2). Mummer4, bowtie2 and blastn all maintain a detection rate close to bowtie1. For low-occurrence spacers (1-10 occurrences), strobealign achieves detection rates of 95.44% but shows a systematic decline to approximately 20% for spacers occurring >100 times, and further drops below 5% in the high occurrence range (>1000).

**Figure 2:**
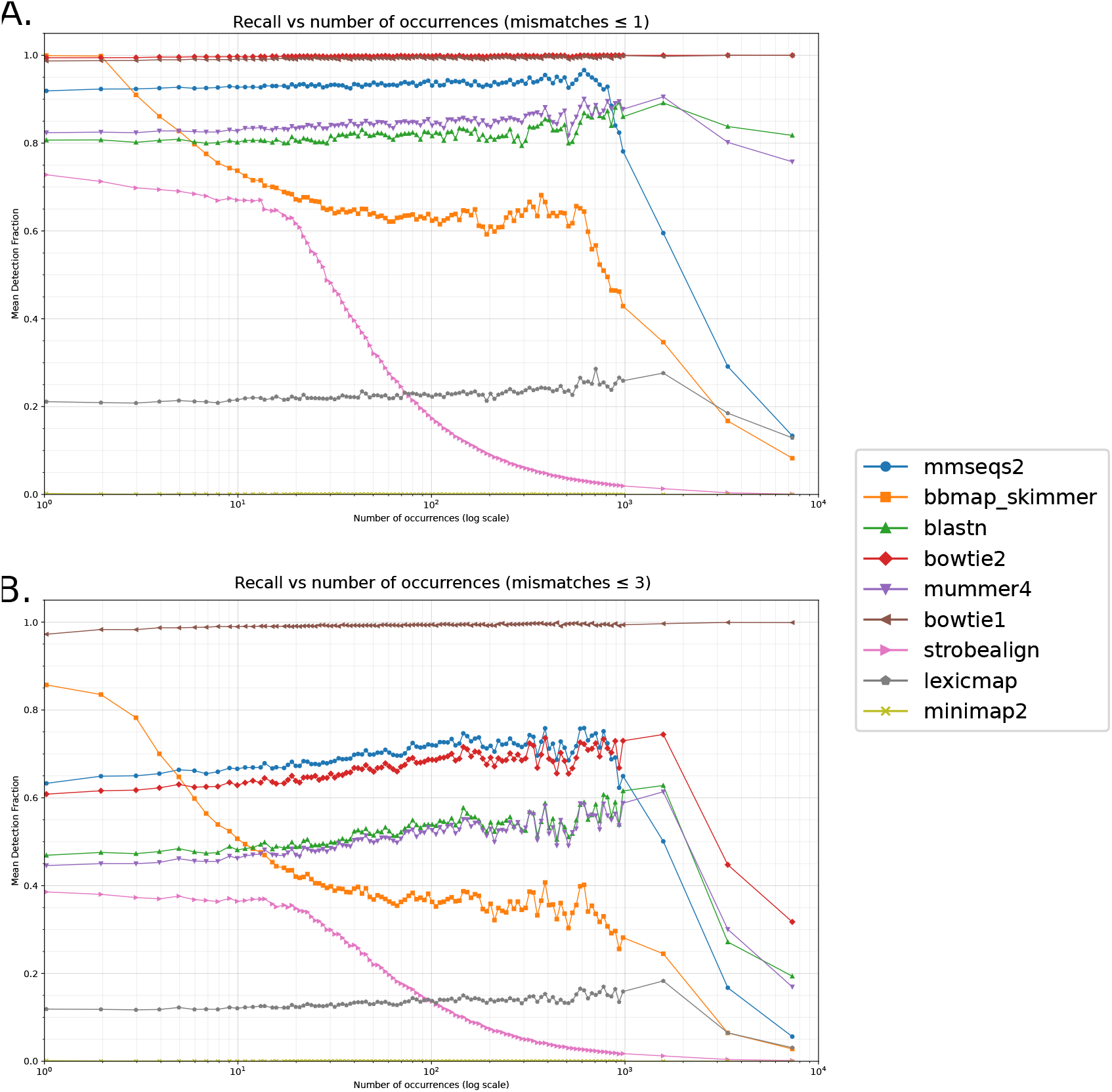
Comparison of recall (detection rate) across different mismatch thresholds and target abundance levels for IMG/VR v4 virus and spacer dataset. Top panel displays the subset of results with up to 1 mismatch, and the bottom panel displays the results with up to 3 mismatches. The horizontal axis shows the number of target occurrences on a logarithmic scale from 1 to 10^4^, while the vertical axis represents the mean detection fraction (0-1). Each color and shape indicates a different tool plot (shapes connected by lines for interpolation). The low-abundance region (1 - 1000 occurrences) is binned into logarithmically-spaced bins, while the high-abundance region (>1000 occurrences) is divided into only 3 additional bins, as such ultra-high abundance sequences are rare. The detection fraction is the mean detection fraction across all spacer-contig pairs at a given mismatch threshold and target abundance level.

When allowing one mismatch, the overall detection capabilities decrease across all tools, although Bowtie1 maintains its high performance. At up to three mismatches, the overall recall rates for all other tools further decrease, while Bowtie1 maintains detection rates above 97% throughout the occurrence spectrum.

The data shows a consistent pattern where detection rates generally decline for spacers with very high occurrence frequencies (>1000), though this effect becomes less pronounced as more mismatches are permitted. Quantitatively, this decline is most evident in tools like strobealign and bbmap-skimmer, while bowtie1 maintains its high performance even with highly repetitive sequences. Detailed statistics and recall curves for exact mismatch values (rather than at a maximal value) can be found in the supplementary.

### 4.3 Overall number of identified spacer-contig

The pairwise comparison of the tool results suggests (Figure 3), reinforces the observation regarding bowtie1’s unique ability to recover a maximum number of spacer matches. Generally, it appears that, when compared to any single other tool, the total number of contig-spacer pairs bowtie1 misses is relatively smaller than the number of pairs the compared tool identified which were not identified by bowtie1.

**Figure 3:**
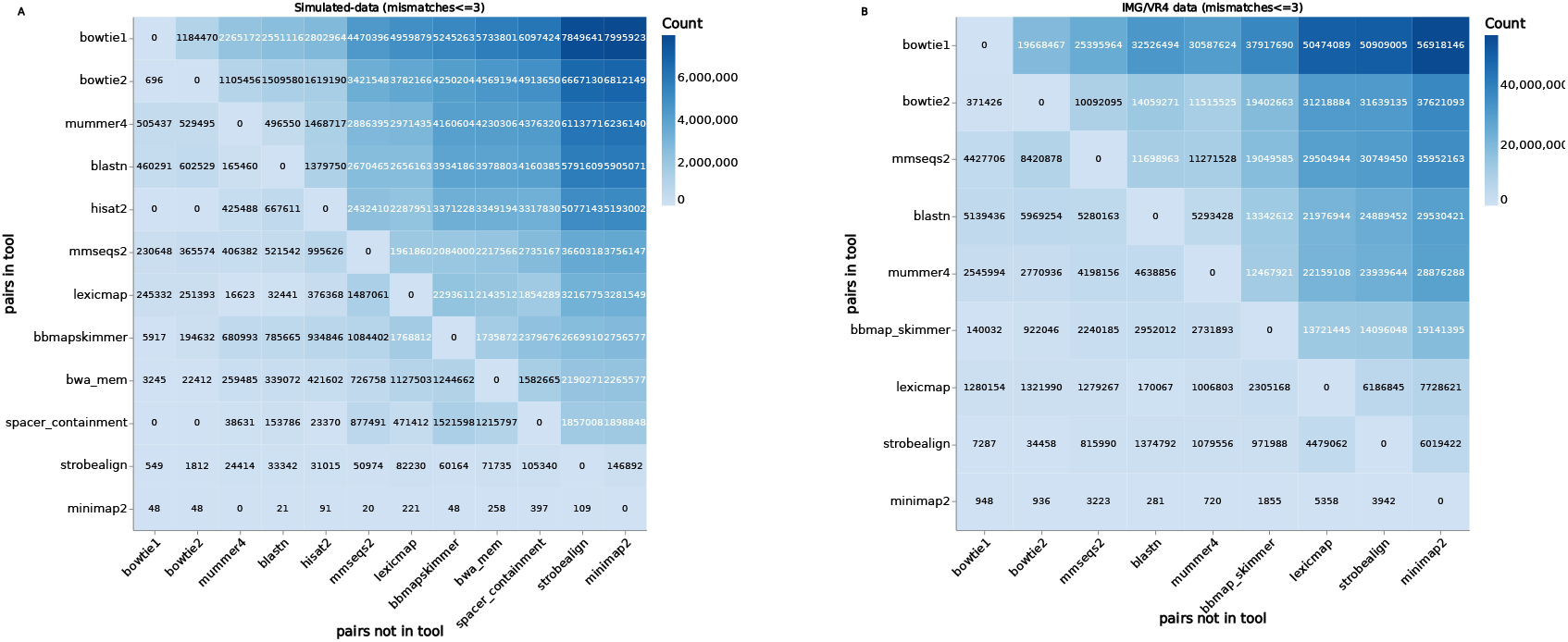
Tool vs Tool (pairwise) comparisons - set intersections and differences matrixes. The value of a cell(i,j) is number of spacer-contig pairs identified by the tool listed in row i, which were not identified by the tool listed in the j column. Panel A shows the results for the synthetic dataset, while panel B shows the results for the IMG/VR4 dataset.

## 5 Discussion

Our findings reveal that within the mismatch thresholds tested (less or equal to 3), no single tool was able to identify all spacer occurrences. While we observe variations across tools, the main difference appears to be the effect of the abundance of the spacer in the target set on the recall rate. Of specific concern is the relatively high number of alignments missed by blastn-short, which is the common method currently used.

In this context, missing potential spacer-protospacer pairs can significantly impact downstream conclusions, e.g. about host range of a given MGE. Overall, Bowtie1 presented the highest recall, although still imperfect for non exact matches. It should be noted that bowtie1 is not designed to identify >3 mismatches, and so a truly exhaustive approach might be to utilize a combination of tools to increase the recall rate, and use a post-search verification method (such as the score recalculation performed here using parasail) to remove potential false positives. Potentially, low complexity sequence filtering (or masking) could be performed prior to the search.

It should be noted that experimental and analytical context may have to be considered when deciding on a specific tool, especially in settings where a higher level of sequence dissimilarity could be considered. On one hand, spacer and virus databases can be expected to continue to increase in size, and our results suggest that large-scale meta-analysis studies (in which spacers from multiple samples are compared to a many potential targets that may include similar or identical regions), should favour tools with high recall, to circumvent potential reporting limitations other tools might have. This is especially important as in such meta-analysis, it should not be assumed the spacer and the potential target co-occur (from the same biological sample) and though the similarity may imply a past encounter between the ancestors of the sequences, caution is due in inferring that the current MGE sequence is capable of infecting the host sequence. In contrast, in experimental designs where potential CRISPR-target pairs are constrained by prior knowledge, tools with higher tolerance for sequence mismatches such as bowtie2 or BLASTn may be more appropriate (e.g. when studying known phage-host pairs from isolates, or when investigating temporally resolved samples (e.g. time-series)).

Another consideration should be the source of the spacer data: spacers sequences extracted from raw NGS data and spacers extracted from assembled CRISPR arrays (either from assembled or long read sequencing). Specifically, spacers from complete arrays present additional information, namely the location and order of the spacers within the array, and the observation they originate from the same host genome. Notably, a recent in-depth study by Mitrofanov et al Mitrofanov et al. [18], investigating the mutational landscape of repeats across many isolate prokaryote genomes, have identified patterns of spacer loss based on system sub/type and location. A similar meta-analysis of spacer mutations by Vink et al Vink, Baijens, and Brouns [32], have revealed that different CRISPR subtypes exhibit varying tolerance for mismatches within the spacer sequences, with most matched spacers containing three or fewer mismatched nucleotides. This aligns with our current general recommendation of using Bowtie1. Additionally, Vink et al observed that Type I-E and Type II systems preferentially target template strands while Type I-A, I-B, and Type III systems prefer coding strands, emphasizing system-specific characteristics which may also inform post-search verification methods (albeit this may require additional information, such as the sequences orientation or coding potential, and the CRISPR subtype of the spacer).

While our technical comparison focuses on tool performance, the biological interpretation of identified matches also requires careful consideration. The arms race between prokaryotes and MGEs creates a complex landscape where simple sequence matching may not directly translate to genuine host-parasite relationships. We identified several scenarios that could lead to false positive assignments:

### 1. Low Complexity Sequence Matches

Independently of the tool choice, low-complexity (regions with highly skewed GC content, or composed of many repeated sequences) can be susceptible to spurious matches. Low complexity regions may be present in both the virus target set, or in the spacer set, where some non-CRISPR repeated sequences may have been misclassified as such. While certain tools employ filters and heuristics to mitigate the effect of low complexity regions, a prudent procedure should include a step separated from the search, to specifically identify, filter or mask the spacer and virus sets. Dustmasker [20], or a similar tool (e.g. ldust from minimap, or BBDuk from bbmap) could be used prior to the search.

### 2. Common Sequence Motifs

Some matches may correspond to highly conserved sequences shared across various biological systems. For instance, horizontal gene transfer events can lead to the spread of similar sequences across diverse MGEs, potentially creating spurious matches that don’t reflect direct host-MGE interactions. An example of potential HGT mediated matches was described by Kosmopoulos et al. 2023, where a transposon-mediated transferred of a phage lysin gene (to the host genome) created a true sequence similarity (which was verified by the authors using a combination of sequencing technologies) [12]. Even if anecdotally observed, this suggests that an unknown number of observed “good” alignments may be due to HGT, which in the absence of additional information, could not be ruled out as a false positive.

#### 3. Self-Targeting Events

Some matches may represent CRISPR targeting of host genes [33]. Previous studies estimated a varying amount of these actually target sequences with putative exogenous origin such as prophages, ranging from ∼50% [30], to ∼80% [28]. In Shmakov et al., the authors estimate non-defence targeting is likely a rare event. So far, most observations of non-defence (or counter defence) molecular functions of CRISPRs did not directly involve the spacer sequences, but rather the Cas genes or related effectors. Some observed functions include genome remodelling and evolution, or temporal regulation of gene expression. For example, in *Francisella novicida*, Sampson et al. 2013 demonstrated that certain lipoprotein production is mediated by a CRISPR system [25].

#### 4. non-chromosomal replicon encoded CRISPRs

Similarly to the potential of CRISPR systems to act in non-immune functions, in certain scenarios spacers may be acquired from non-MGE replicons, or be carried (even if partially) by mobile elements. While some types are known to be chromosomal, others are known to be carried entirely by plasmids (i.e. both the Cas proteins and the array loci are on the plasmid), for example in various halophilic archaea [15]. A recent study by Zhang et al [34] observed similar phenomena in the human gut microbiome, specifically in *Bifidobacterium longum*. Another non-MGE targeting phenomena in archaea was described by Turgeman-Grott et al. [31], where inter-species spacers (targeting genes from related species) were demonstrated to be common in archaea, at least in the context of cellular mating. Another confounding factor is the potential of certain MGEs to target host genes, potentially for regulatory functions, or as counter-defense mechanisms[27]. Shmakov et al. 2023 have identified widespread CRISPR-derived phage-encoded mini-arrays, which can hijack and interfere with their host native system.

#### 5.1 Study Limitations

The synthetic data was generated with uniform distributions of spacer and contig lengths and base composition (where each nucleotide is chosen independently and at random). While the simulation scripts do allow user-specified base composition (or use of real data via the –contigs or –spacers parameters), real biological sequence composition is not usually uniformly distributed, and may be biased towards certain nucleotide frequencies, which could be region or loci specific. This can be clearly observed comparing the spacer attributes distributions in the synthetic and real datasets (see supplementary figure 6), and table 1. While the simulated dataset sample size is relatively small, the high level of control over the spacer and contig sequences allowed testing rare or yet-unobserved scenarios, like ultra high spacer occurrence rates. This difference in sample size and features should be considered when interpreting the results.

Furthermore, we focused on a specific distance metric (Levenshtein distance), thus the synthetic sequences only have substitutions simulated into them. While insertions and deletions were not simulated, and although these might be relatively rare in short evolutionary time, they are still possible and could inform real and recent MGE-host interactions (e.g an indel that does not disrupt the encoded sequence reading frame). This could be further explored in future work, and might affect overall tool suggestion to use bowtie1 for general-case spacer-protospacer matching, as it is not designed with gaps in mind. Finally, we note that tools were not tested for all possible combinations of their available tunable parameters, and it is possible some might be further tweaked to improve their performance. Similarly, variation in parameters during index construction may further affect this (e.g. offset rate, seed selection). Similarly, not all major versions of all tools were tested. Of specific note are MMseqs2 (where the latest release version at the time of execution resulted in frequent crashes, hence we used the latest commit version from github), and Lexicmap which does not have a formal “v1” release at the time of writing, and since the benchmark was run, a new minor version was released, which is a good candidate for future testing.

## 6 Conclusion

Our comparison of spacer-protospacer search tools reveals variations in their ability to detect unique putative CRISPR-target matches, which is a key and common input for current host-MGE interaction tools and studies. The main tool-derived difference we report is the ability to detect multiple matches - most tools show a sharp decline in detection rates as the total number of occurrences increases. This can directly impact the downstream interpretation of microbial relationships, such as under estimation of potential hosts-range, or over reliance on seemingly unique matches which may actually be due to non-direct intercations resulting from common sequence motifs or high tolerance for mismatches. While Bowtie1 demonstrates superior performance in detecting multiple matches under the threshold of 3 mismatches, the interpretation of spacer-protospacer matches requires careful consideration of biological context and potential interfering factors, especially when comparing large databases of CRISPR spacers and potential targets collected from a broad range of samples and ecosystems. Our findings also emphasize the importance of recalculating alignment metrics in a tool-agnostic manner, to ensure consistent comparison across tools. Finally, we provide general recommendations and experimental design considerations of potential interfering factors for future studies.

## 7 Code and data availability

All code generated for this study can be found in the git repository: code.jgi.doe.gov/spacersdb/spacer_matching_bench. All raw outputs (tool results on real and synthetic datasets, the simulated sequence files, the SLURM logs, and the hyperfine runtime measurements) are available on Zenodo [21].

## 8 Acknowledgements

Work conducted by the U.S. DOE Joint Genome Institute (https://ror.org/04xm1d337) (SR, UN, APC and BB), a DOE Office of Science User Facility, is supported by the Office of Science of the U.S. DOE operated under Contract DE-AC02-05CH11231.

We would like to thank the following people for their helpful feedback and suggestions: Uri Gophna and Georg Rath.

## References

[1] S F Altschul et al. “Basic local alignment search tool”. In: J. Mol. Biol. 215.3 (Oct. 1990), pp. 403–410.

[2] Ambarish Biswas et al. “CRISPRTarget”. In: RNA Biology 10.5 (2013), pp. 817–827. doi: 10.4161/rna.24046.

[3] Charles Bland et al. “CRISPR Recognition Tool (CRT): a tool for automatic detection of clustered regularly interspaced palindromic repeats”. In: BMC Bioinformatics 8.1 (Dec. 2007), p. 209. issn: 1471-2105. doi: 10.1186/1471-2105-8-209. url: https://bmcbioinformatics.biomedcentral.com/articles/10.1186/1471-2105-8-209.

[4] Antonio Pedro Camargo et al. “Identification of mobile genetic elements with geNomad”. en. In: Nature Biotechnology 42.8 (Aug. 2024), pp. 1303–1312. issn: 1087-0156, 1546–1696. doi: 10.1038/s41587-023-01953-y. url: https://www.nature.com/articles/s41587-023-01953-y.

[5] Antonio Pedro Camargo et al. “IMG/VR v4: an expanded database of uncultivated virus genomes within a framework of extensive functional, taxonomic, and ecological metadata”. In: Nucleic Acids Research 51.D1 (Jan. 2023), pp. D733–D743. issn: 0305-1048, 1362-4962. doi: 10.1093/nar/gkac1037. url: https://academic.oup.com/nar/article/51/D1/D733/6833254.

[6] conda contributors. conda: A system-level, binary package and environment manager running on all major operating systems and platforms. url: https://github.com/conda/conda.

[7] Jeff Daily. “Parasail: SIMD C library for global, semi-global, and local pairwise sequence alignments”. In: BMC Bioinformatics 17.1 (Feb. 2016), p. 81. issn: 1471-2105. doi: 10.1186/s12859-016-0930-z. url: https://doi.org/10.1186/s12859-016-0930-z.

[8] Moïra B Dion et al. “Streamlining CRISPR spacer-based bacterial host predictions to decipher the viral dark matter”. In: Nucleic Acids Research 49.6 (Mar. 2021), pp. 3127–3138. issn: 1362-4962. doi: 10.1093/nar/gkab133. url: http://dx.doi.org/10.1093/nar/gkab133.

[9] Robert C Edgar. “PILER-CR: Fast and accurate identification of CRISPR repeats”. In: BMC Bioinformatics 8.1 (Dec. 2007), p. 18. issn: 1471-2105. doi: 10.1186/1471-2105-8-18. url: https://bmcbioinformatics.biomedcentral.com/articles/10.1186/1471-2105-8-18.

[10] Robert A. Edwards et al. “Computational approaches to predict bacteriophage–host relationships”. In: FEMS Microbiology Reviews 40.2 (Dec. 2015), pp. 258–272. issn: 0168-6445. doi: 10.1093/femsre/fuv048. eprint: https://academic.oup.com/femsre/article-pdf/40/2/258/23905864/fuv048.pdf. url: https://doi.org/10.1093/femsre/fuv048.

[11] Eugene V. Koonin and Kira S. Makarova. “Evolutionary plasticity and functional versatility of CRISPR systems”. In: PLOS Biology 20.1 (Jan. 2022), pp. 1–19. doi: 10.1371/journal.pbio.3001481. url: https://doi.org/10.1371/journal.pbio.3001481.

[12] James C. Kosmopoulos et al. “Horizontal Gene Transfer and CRISPR Targeting Drive Phage-Bacterial Host Interactions and Coevolution in “Pink Berry” Marine Microbial Aggregates”. In: Applied and Environmental Microbiology 89.7 (July 2023). Ed. by Martha Vives. issn: 1098-5336. doi: 10.1128/aem.00177-23. url: http://dx.doi.org/10.1128/aem.00177-23.

[13] Vladimir I Levenshtein. “Binary Codes Capable of Correcting Deletions, Insertions and Reversals”. In: Soviet Physics Doklady 10 (Feb. 1966), p. 707.

[14] Thomas L Madden, Ben Busby, and Jian Ye. “Reply to the paper: Misunderstood parameters of NCBI BLAST impacts the correctness of bioinformatics workflows”. In: Bioinformatics 35.15 (Dec. 2018), pp. 2699–2700. issn: 1367-4803. doi: 10.1093/bioinformatics/bty1026. eprint: https://academic.oup.com/bioinformatics/article-pdf/35/15/2699/50722393/bioinformatics\_35\_15\_2699.pdf. url: https://doi.org/10.1093/bioinformatics/bty1026.

[15] Lisa-Katharina Maier et al. “The nuts and bolts of the Haloferax CRISPR-Cas system I-B”. In: RNA Biology 16.4 (May 2018), pp. 469–480. issn: 1555-8584. doi: 10.1080/15476286.2018.1460994. url: http://dx.doi.org/10.1080/15476286.2018.1460994.

[16] Kira S Makarova et al. “Evolutionary classification of CRISPR–Cas systems: a burst of class 2 and derived variants”. In: Nature Reviews Microbiology 18.2 (2020), pp. 67–83.

[17] mamba-org. Mamba: The Fast Cross-Platform Package Manager. url: https://github.com/mamba-org/mamba.

[18] Alexander Mitrofanov et al. “Comprehensive Analysis of CRISPR Array Repeat Mutations Reveals Subtype-Specific Patterns and Links to Spacer Dynamics”. In: bioRxiv (2025). doi: 10.1101/2025.04.02.646798. eprint: https://www.biorxiv.org/content/early/2025/04/02/2025.04.02.646798.full.pdf. url: https://www.biorxiv.org/content/early/2025/04/02/2025.04.02.646798.

[19] Francisco J.M. Mojica et al. “Intervening Sequences of Regularly Spaced Prokaryotic Repeats Derive from Foreign Genetic Elements”. In: Journal of Molecular Evolution 60.2 (Feb. 2005), pp. 174–182. issn: 1432-1432. doi: 10.1007/s00239-004-0046-3. url: http://dx.doi.org/10.1007/s00239-004-0046-3.

[20] Aleksandr Morgulis et al. “A Fast and Symmetric DUST Implementation to Mask Low-Complexity DNA Sequences”. In: Journal of Computational Biology 13.5 (June 2006), pp. 1028–1040. issn: 1557-8666. doi: 10.1089/cmb.2006.13.1028. url: http://dx.doi.org/10.1089/cmb.2006.13.1028.

[21] U. Neri et al. “Supplementary data for CRISPR spacer-protospacer matching benchmarks [Data set]”. In: (). doi: 10.5281/zenodo.15171878.

[22] Uri Neri. pyseff: A Python script to calculate the efficiency of SLURM jobs. url: https://github.com/UriNeri/pyseff.

[23] David Peter. hyperfine. Version 1.16.1. Mar. 2023. url: https://github.com/sharkdp/hyperfine.

[24] Simon Roux et al. “iPHoP: An integrated machine learning framework to maximize host prediction for metagenome-derived viruses of archaea and bacteria”. In: PLOS Biology 21.4 (Apr. 2023), pp. 1–26. doi: 10.1371/journal.pbio.3002083. url: https://doi.org/10.1371/journal.pbio.3002083.

[25] Timothy R. Sampson et al. “A CRISPR/Cas system mediates bacterial innate immune evasion and virulence”. In: Nature 497.7448 (Apr. 2013), pp. 254–257. issn: 1476-4687. doi: 10.1038/nature12048. url: http://dx.doi.org/10.1038/nature12048.

[26] Nidhi Shah et al. “Misunderstood parameter of NCBI BLAST impacts the correctness of bioinformatics workflows”. In: Bioinformatics 35.9 (Sept. 2018), pp. 1613–1614. issn: 1367-4803. doi: 10.1093/bioinformatics/bty833. eprint: https://academic.oup.com/bioinformatics/article-pdf/35/9/1613/48942014/bioinformatics\_35\_9\_1613.pdf. url: https://doi.org/10.1093/bioinformatics/bty833.

[27] Sergey A Shmakov et al. “Widespread CRISPR-derived RNA regulatory elements in CRISPR-Cas systems”. In: Nucleic Acids Research 51.15 (June 2023), pp. 8150–8168. issn: 1362-4962. doi: 10.1093/nar/gkad495. url: http://dx.doi.org/10.1093/nar/gkad495.

[28] Sergey A. Shmakov et al. “The CRISPR Spacer Space Is Dominated by Sequences from Species-Specific Mobilomes”. In: mBio 8.5 (Nov. 2017). Ed. by Michael S. Gilmore. issn: 2150-7511. doi: 10.1128/mbio.01397-17. url: http://dx.doi.org/10.1128/mbio.01397-17.

[29] Paola Soto-Perez et al. “CRISPR-Cas System of a Prevalent Human Gut Bacterium Reveals Hyper-targeting against Phages in a Human Virome Catalog”. In: Cell Host & Microbe 26.3 (Sept. 2019), 325–335.e5. issn: 19313128. doi: 10.1016/j.chom.2019.08.008. url: https://linkinghub.elsevier.com/retrieve/pii/S1931312819304172.

[30] Adi Stern et al. “Self-targeting by CRISPR: gene regulation or autoimmunity?” In: Trends in Genetics 26.8 (Aug. 2010), pp. 335–340. issn: 0168-9525. doi: 10.1016/j.tig.2010.05.008. url: http://dx.doi.org/10.1016/j.tig.2010.05.008.

[31] Israela Turgeman-Grott et al. “Pervasive acquisition of CRISPR memory driven by inter-species mating of archaea can limit gene transfer and influence speciation”. In: Nature Microbiology 4.1 (Nov. 2018), pp. 177–186. issn: 2058-5276. doi: 10.1038/s41564-018-0302-8. url: http://dx.doi.org/10.1038/s41564-018-0302-8.

[32] Jochem N. A. Vink, Jan-Hendrik L. Baijens, and Stan J. J. Brouns. “PAM-repeat associations and spacer selection preferences in single and co-occurring CRISPR-Cas systems”. In: Genome Biology (2021). doi: 10.1186/s13059-021-02495-9.

[33] Franziska Wimmer and Chase L. Beisel. “CRISPR-Cas Systems and the Paradox of Self-Targeting Spacers”. In: Frontiers in Microbiology 10 (Jan. 2020). issn: 1664-302X. doi: 10.3389/fmicb.2019.03078. url: http://dx.doi.org/10.3389/fmicb.2019.03078.

[34] An-Ni Zhang et al. “CRISPR-Cas spacer acquisition is a rare event in human gut microbiome”. In: Cell Genomics 5.1 (Jan. 2025), p. 100725. issn: 2666-979X. doi: 10.1016/j.xgen.2024.100725. url: http://dx.doi.org/10.1016/j.xgen.2024.100725.

[35] Ruoshi Zhang et al. “SpacePHARER: sensitive identification of phages from CRISPR spacers in prokaryotic hosts”. In: Bioinformatics 37.19 (Apr. 2021). Ed. by Alfonso Valencia, pp. 3364–3366. issn: 1367-4811. doi: 10.1093/bioinformatics/btab222. url: http://dx.doi.org/10.1093/bioinformatics/btab222.

