## Supplementary figures and images for "Tool Choice drastically Impacts CRISPR Spacer-Protospacer Detection"

### Supplementary figure 1

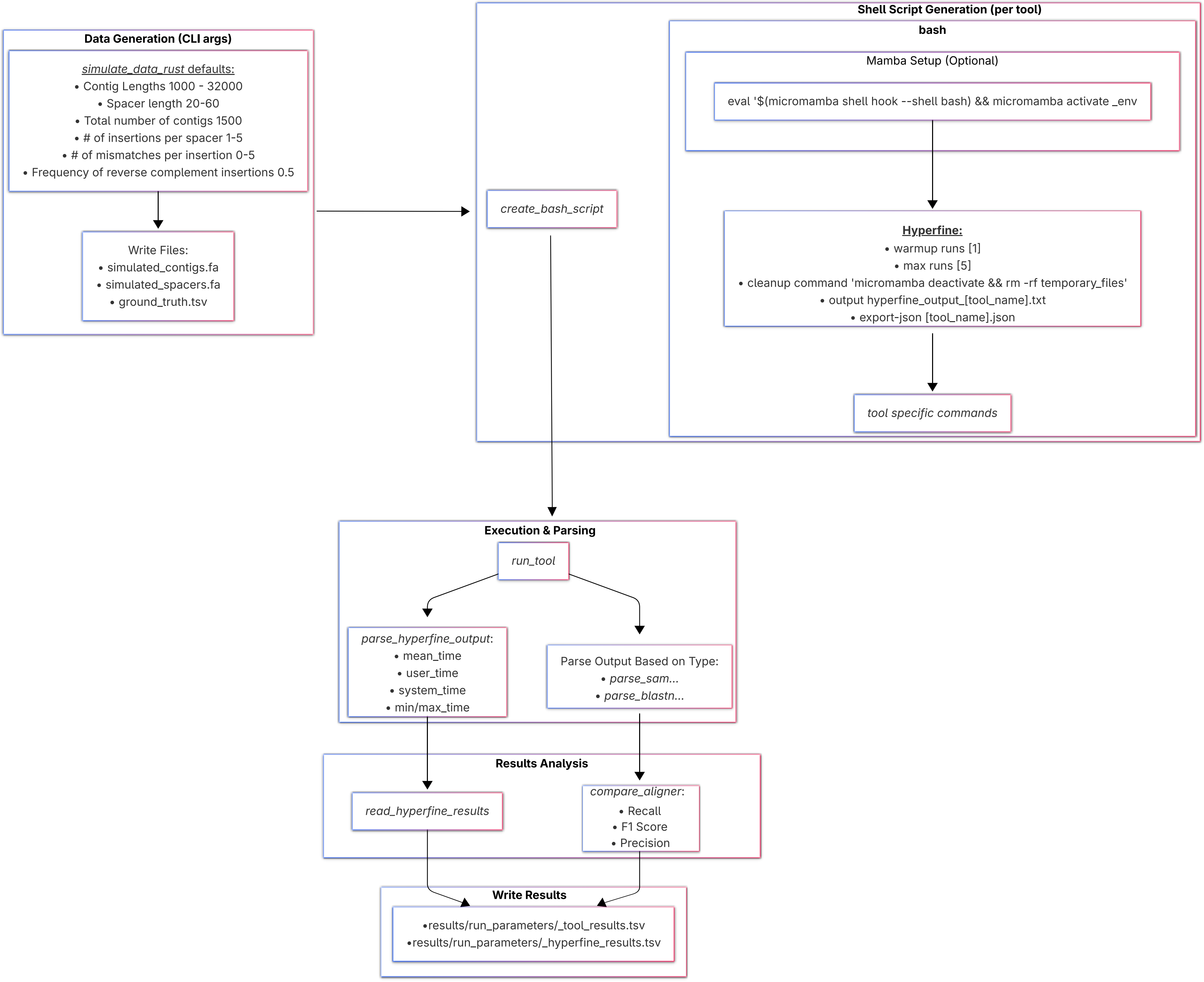

### Supplementary figure 2

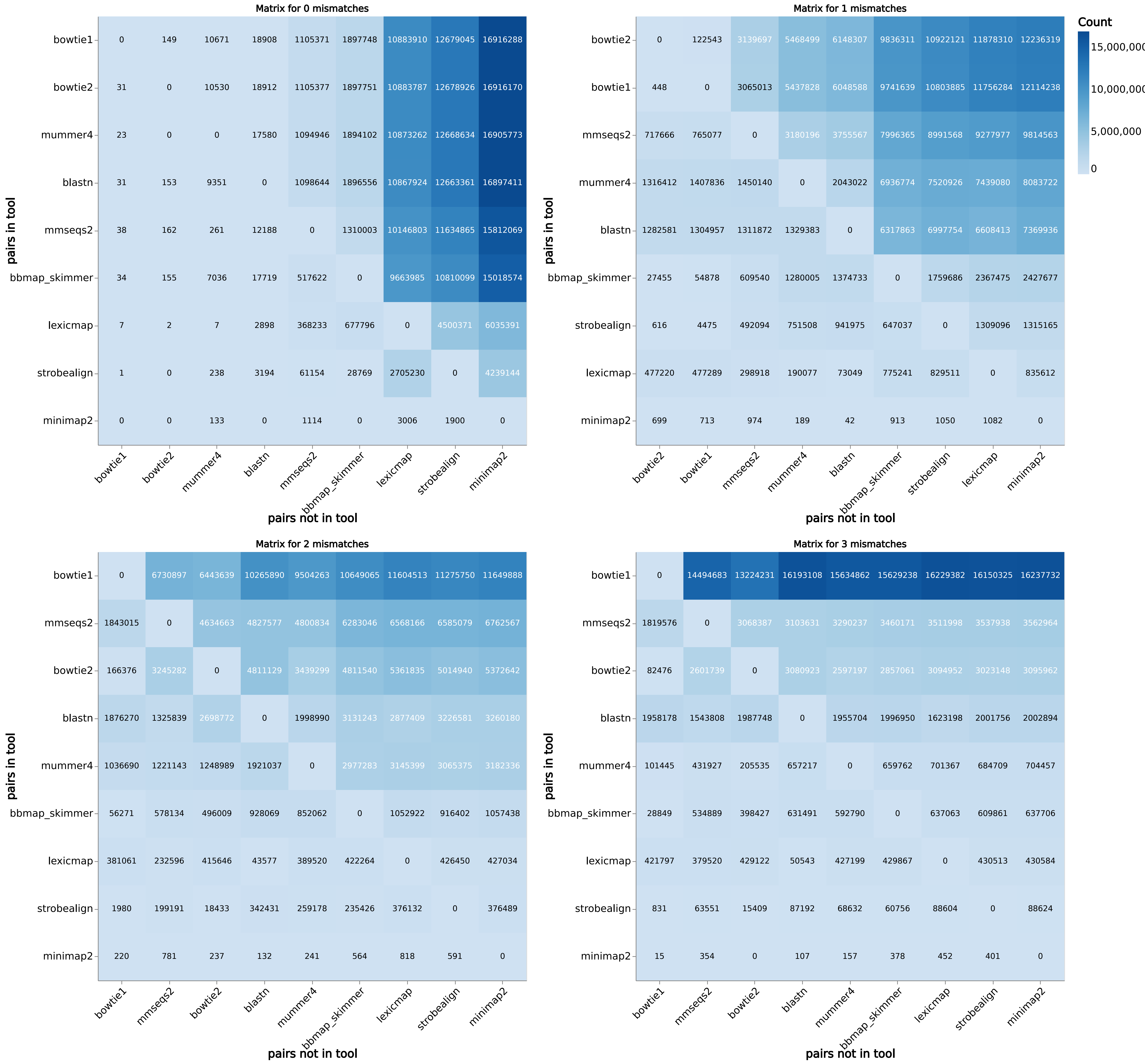

### Supplementary figure 3

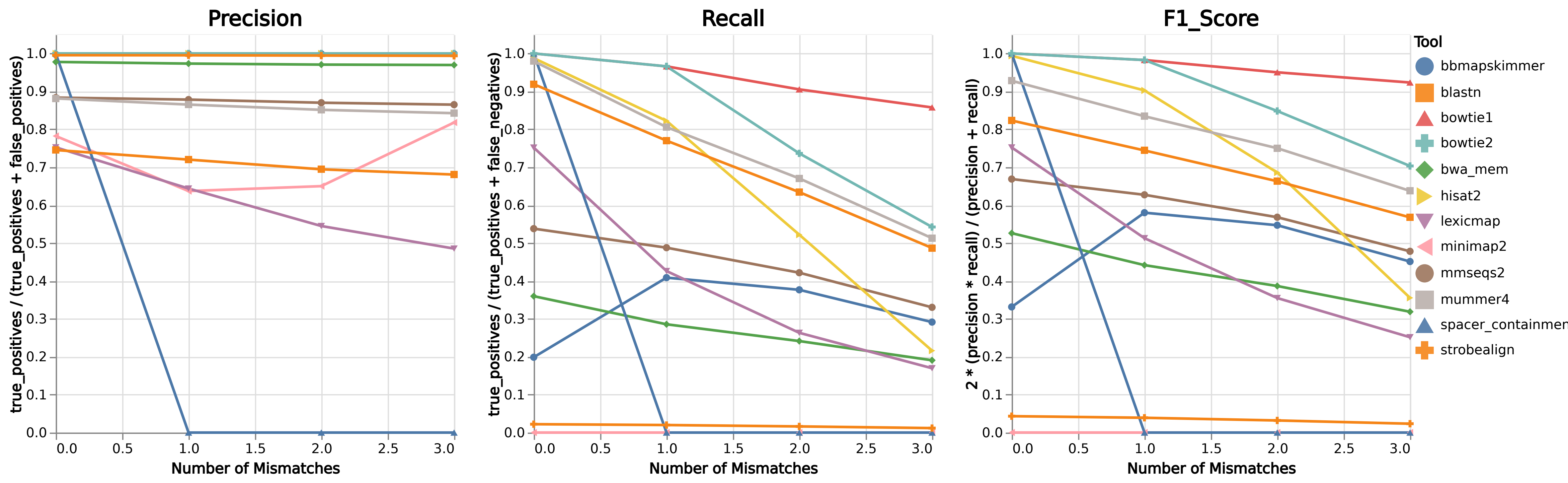

### Supplementary figure 4.A.1.

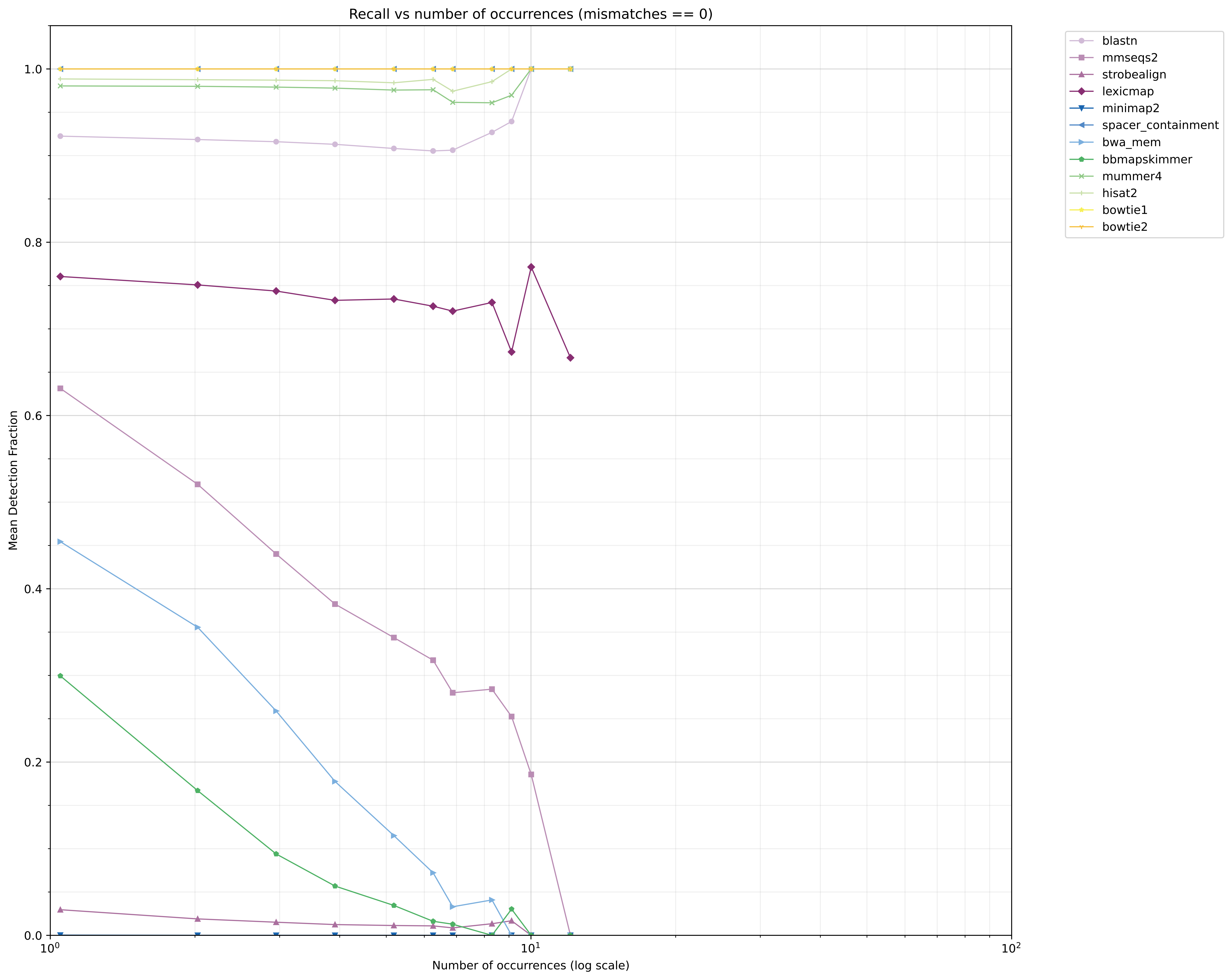

### Supplementary figure 4.A.2.

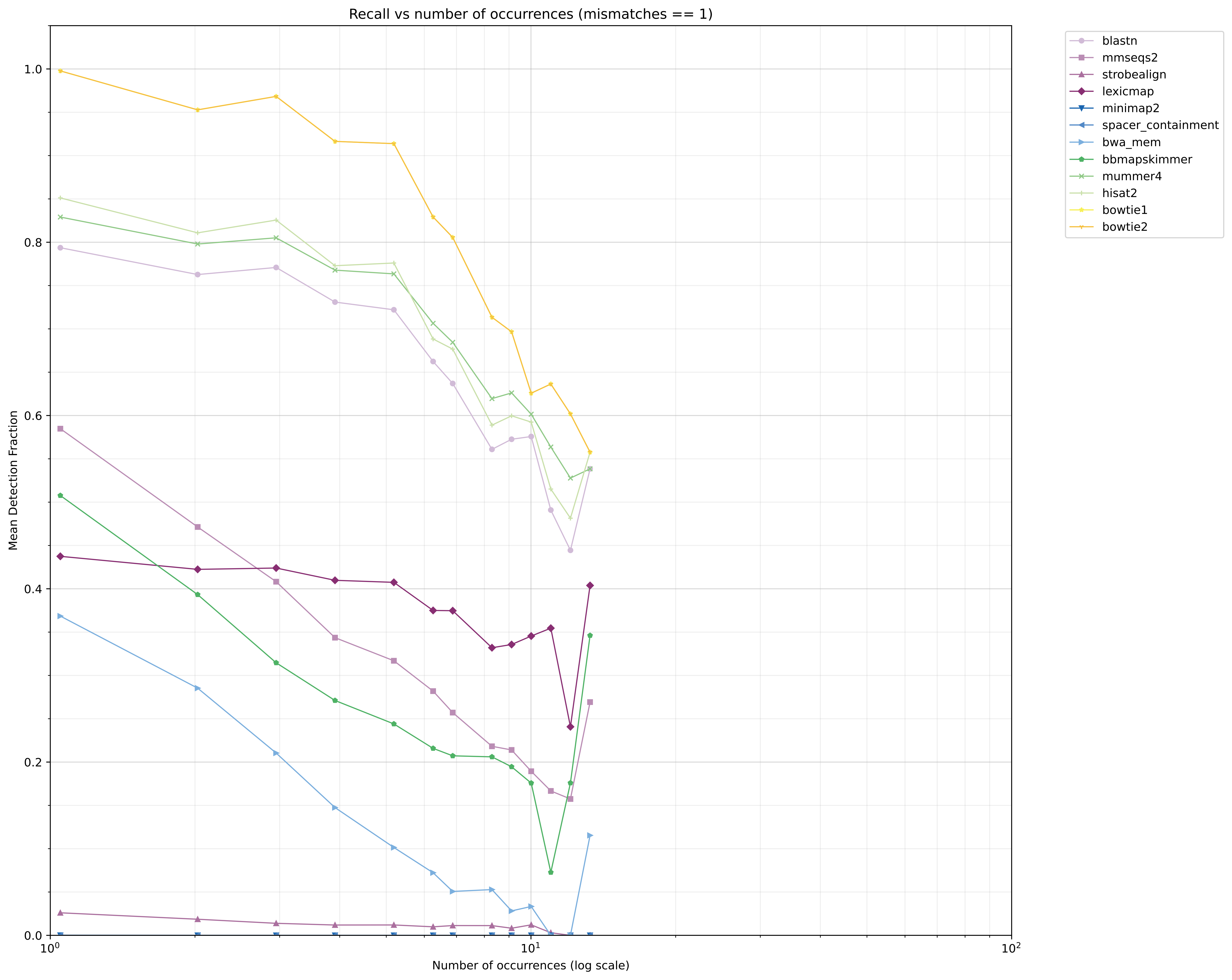

### Supplementary figure 4.A.3.

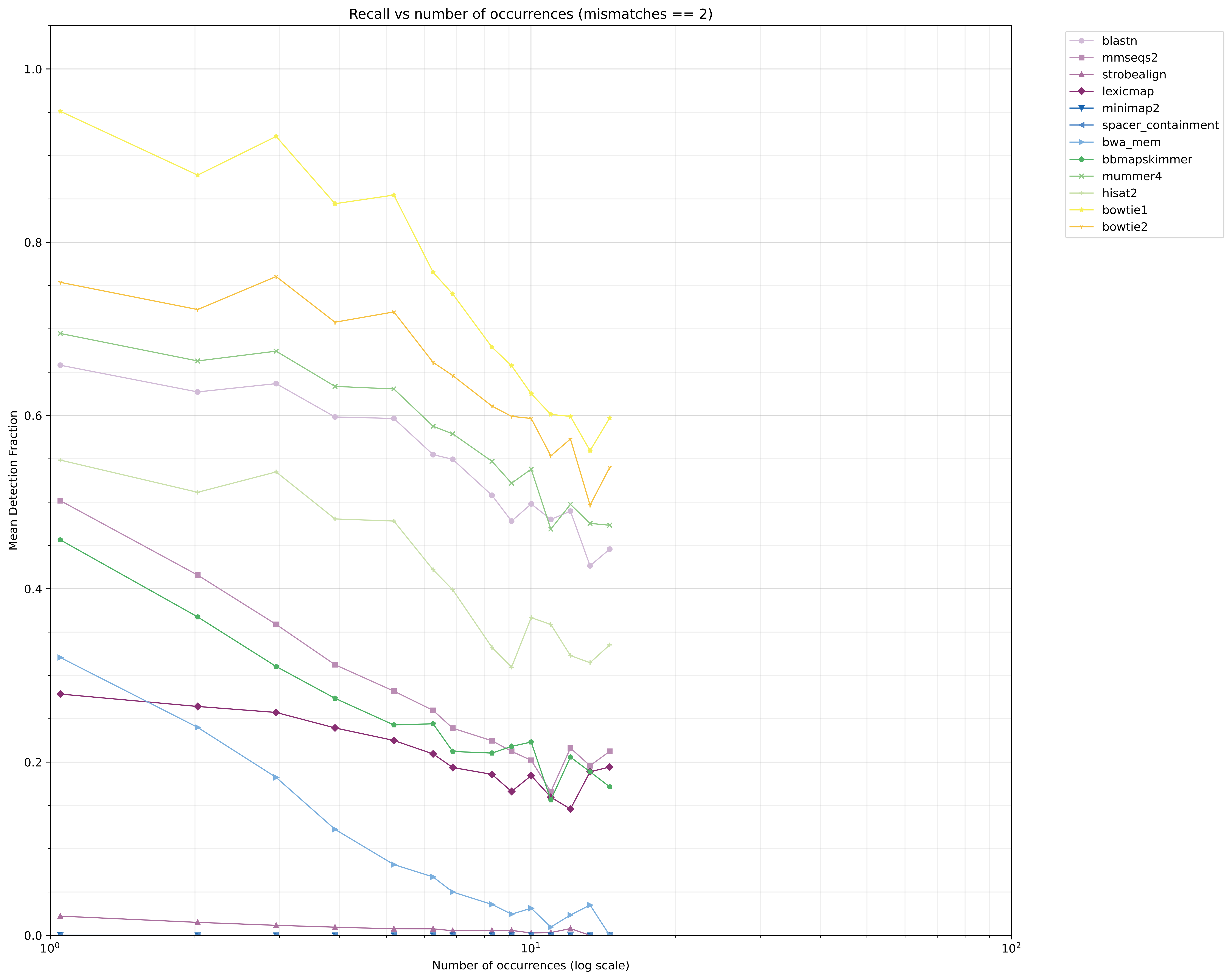

### Supplementary figure 4.A.4.

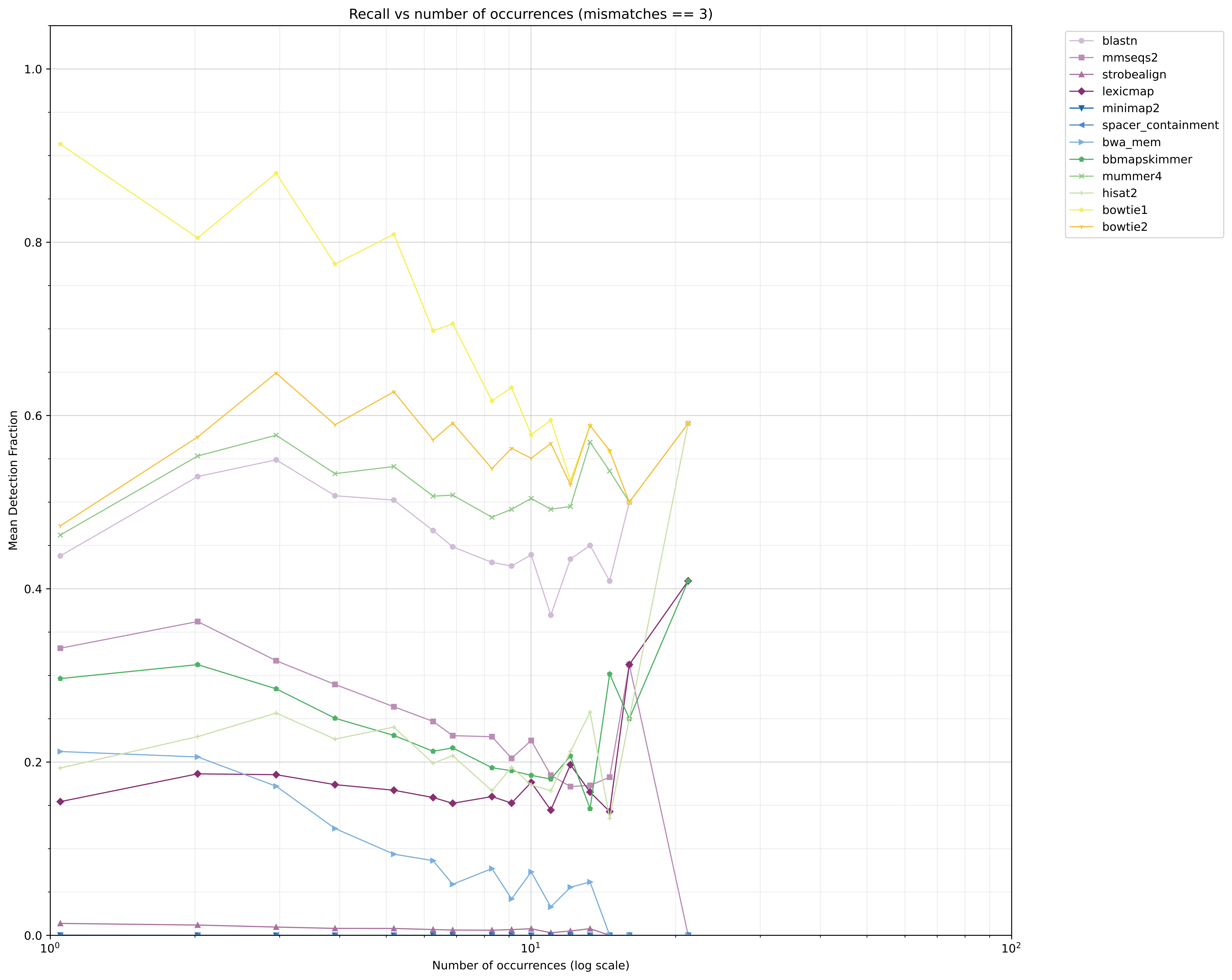

### Supplementary figure 4.B.1.

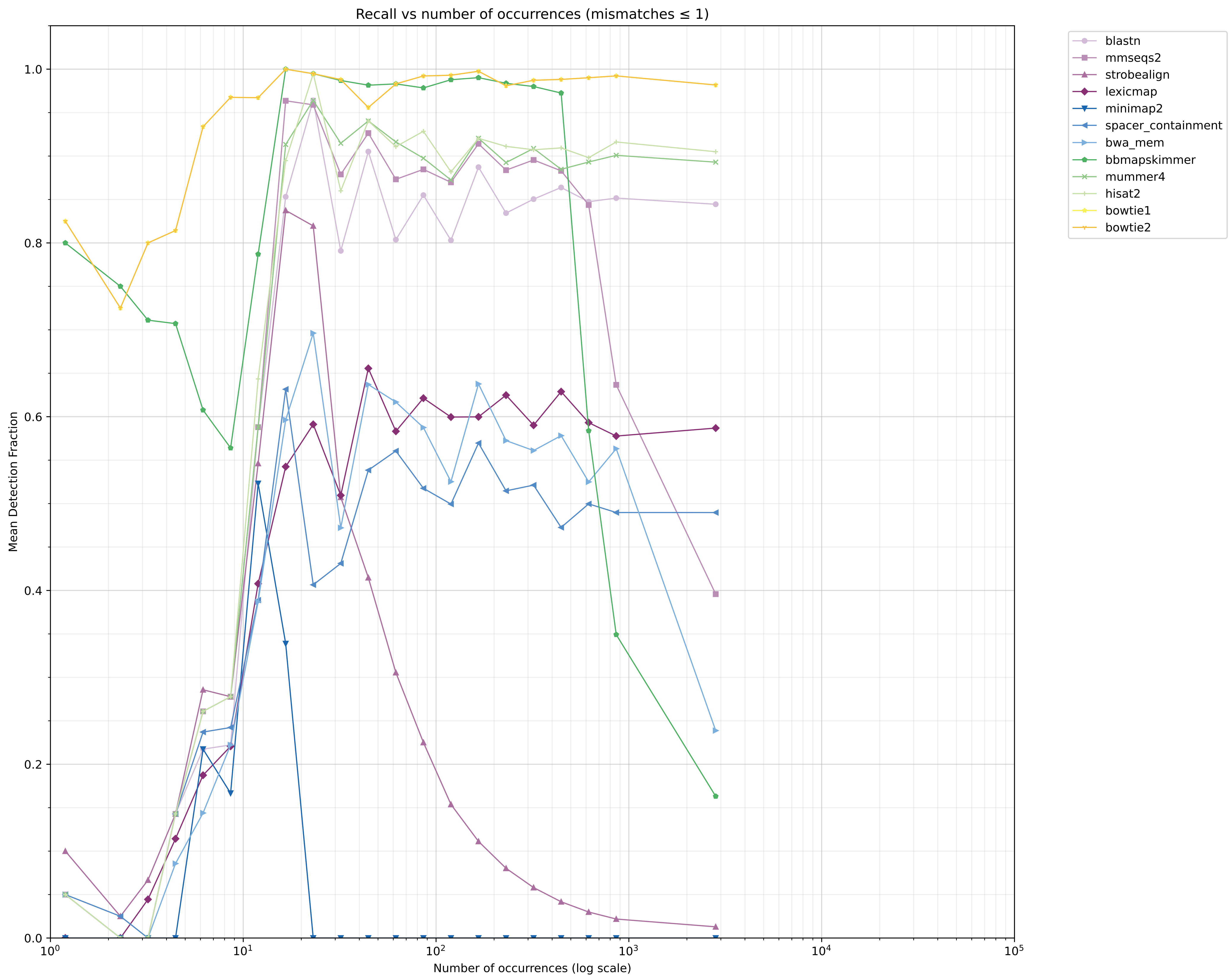

### Supplementary figure 4.B.2.

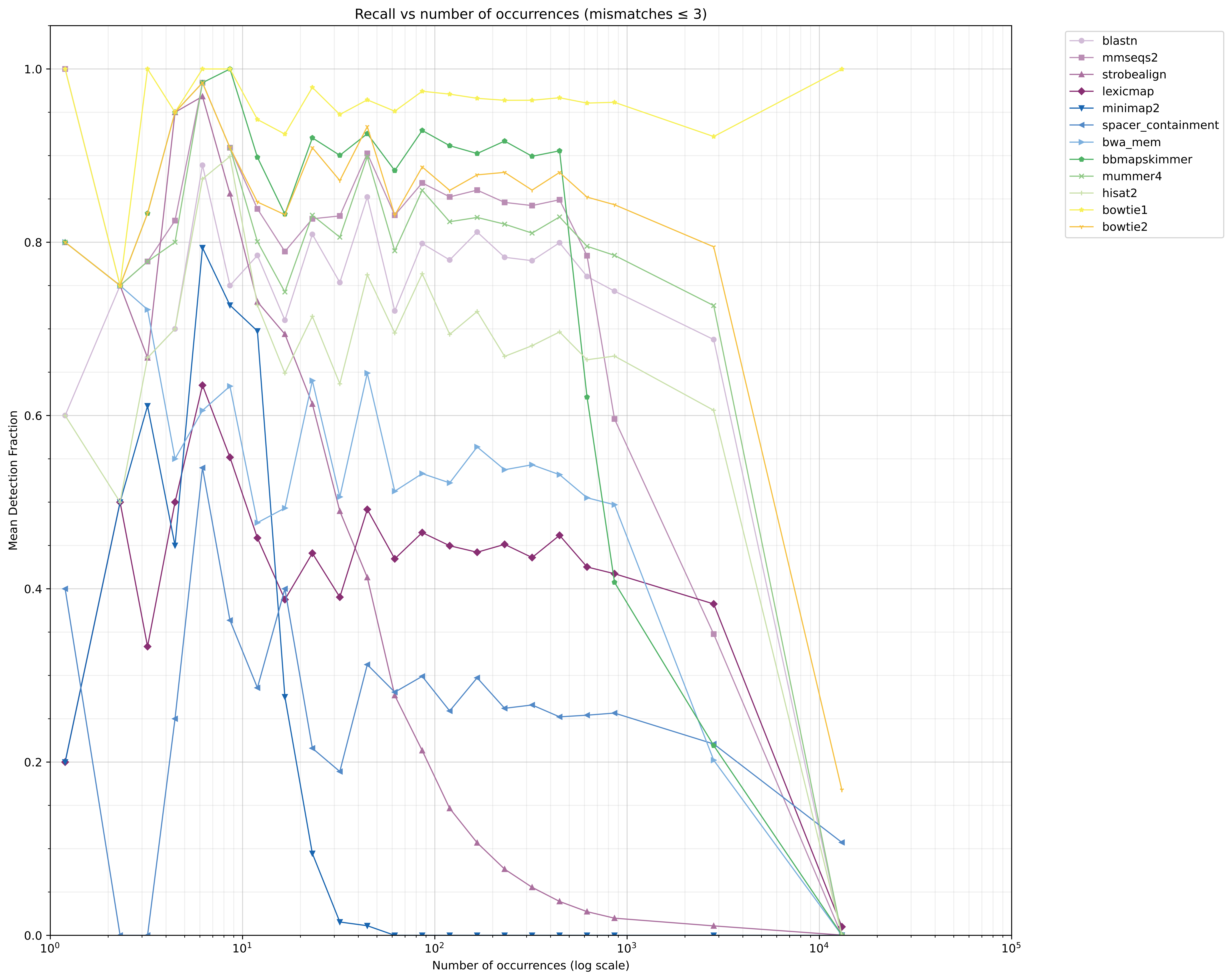

### Supplementary figure 5.A.

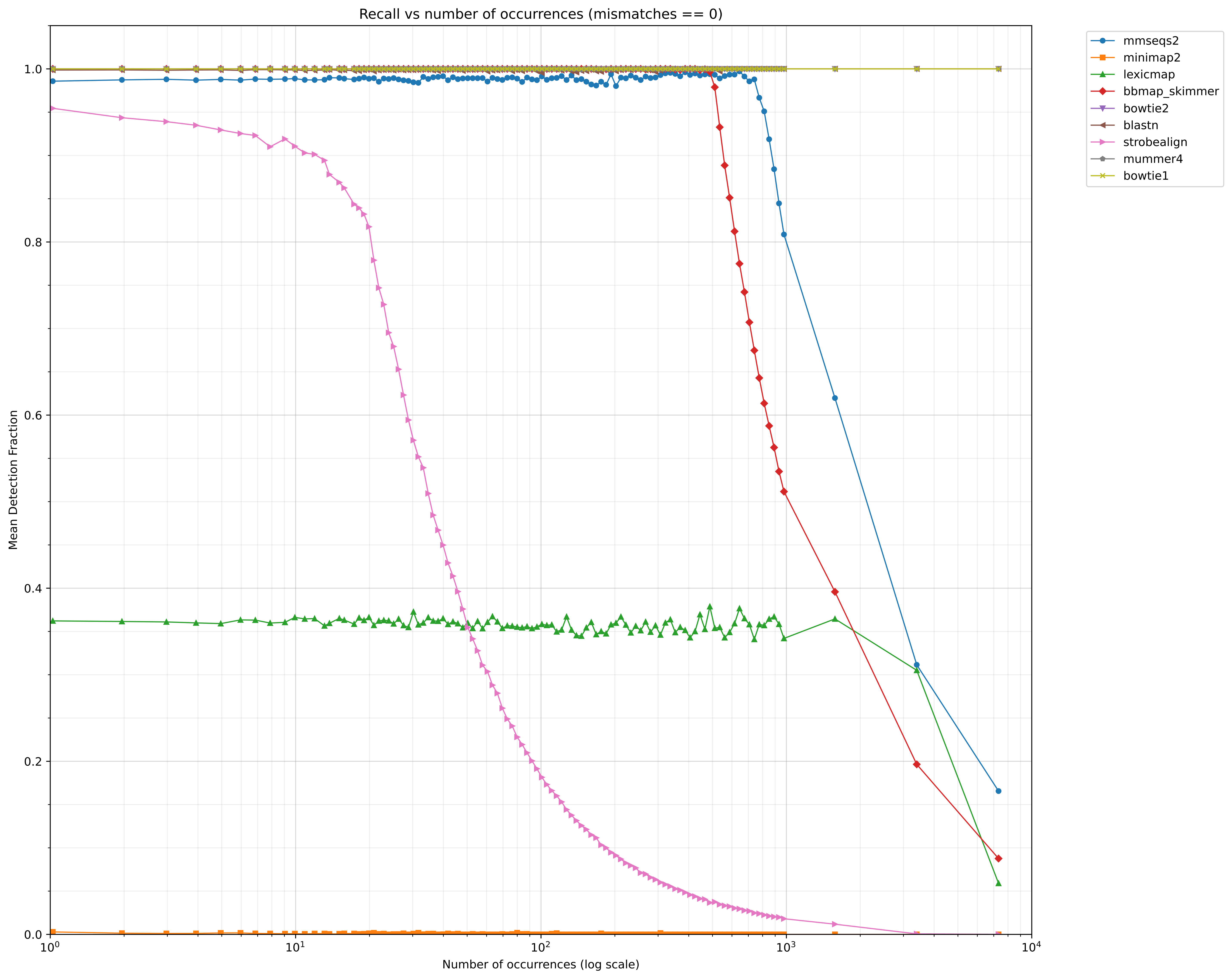

### Supplementary figure 5.B.

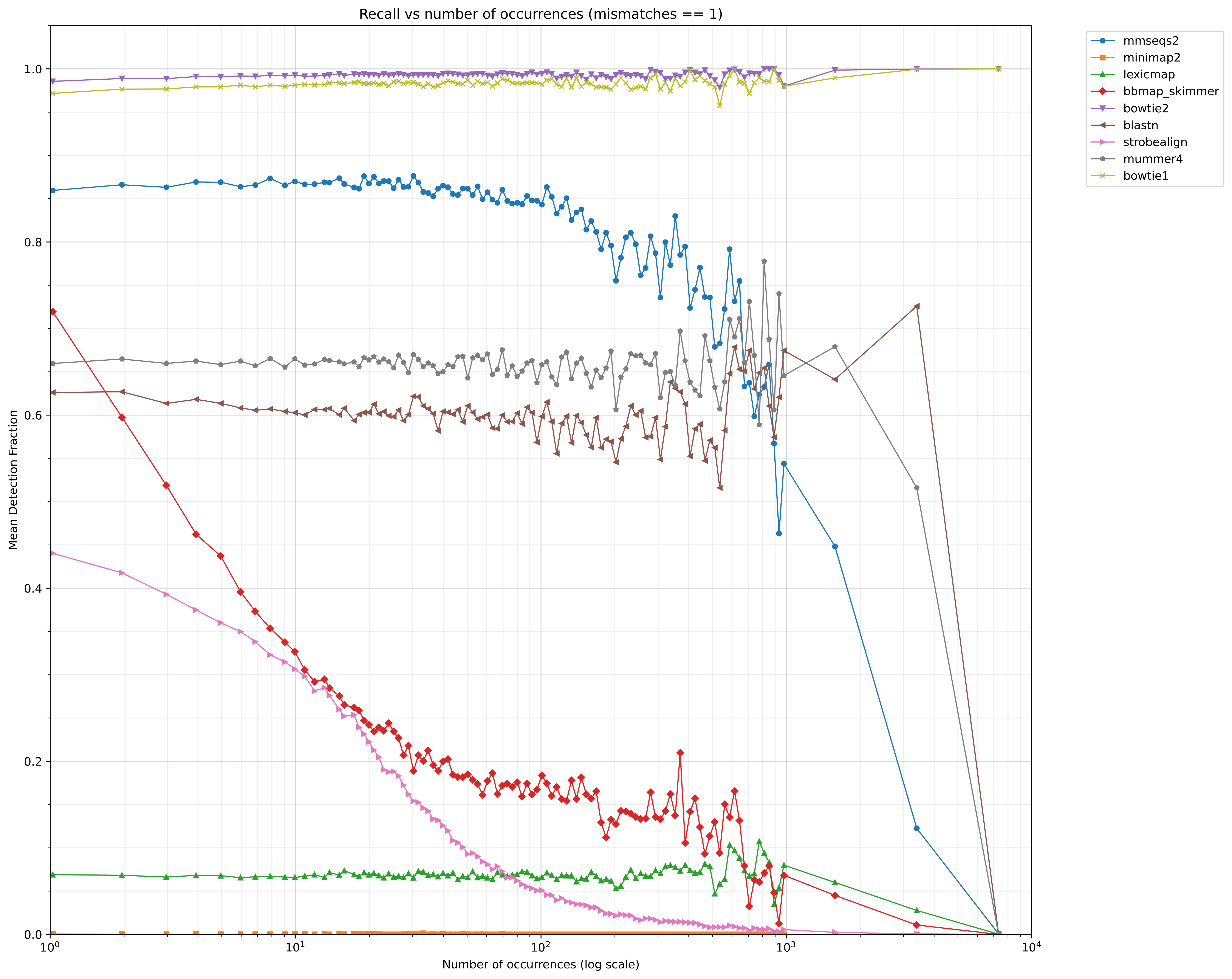

### Supplementary figure 5.C.

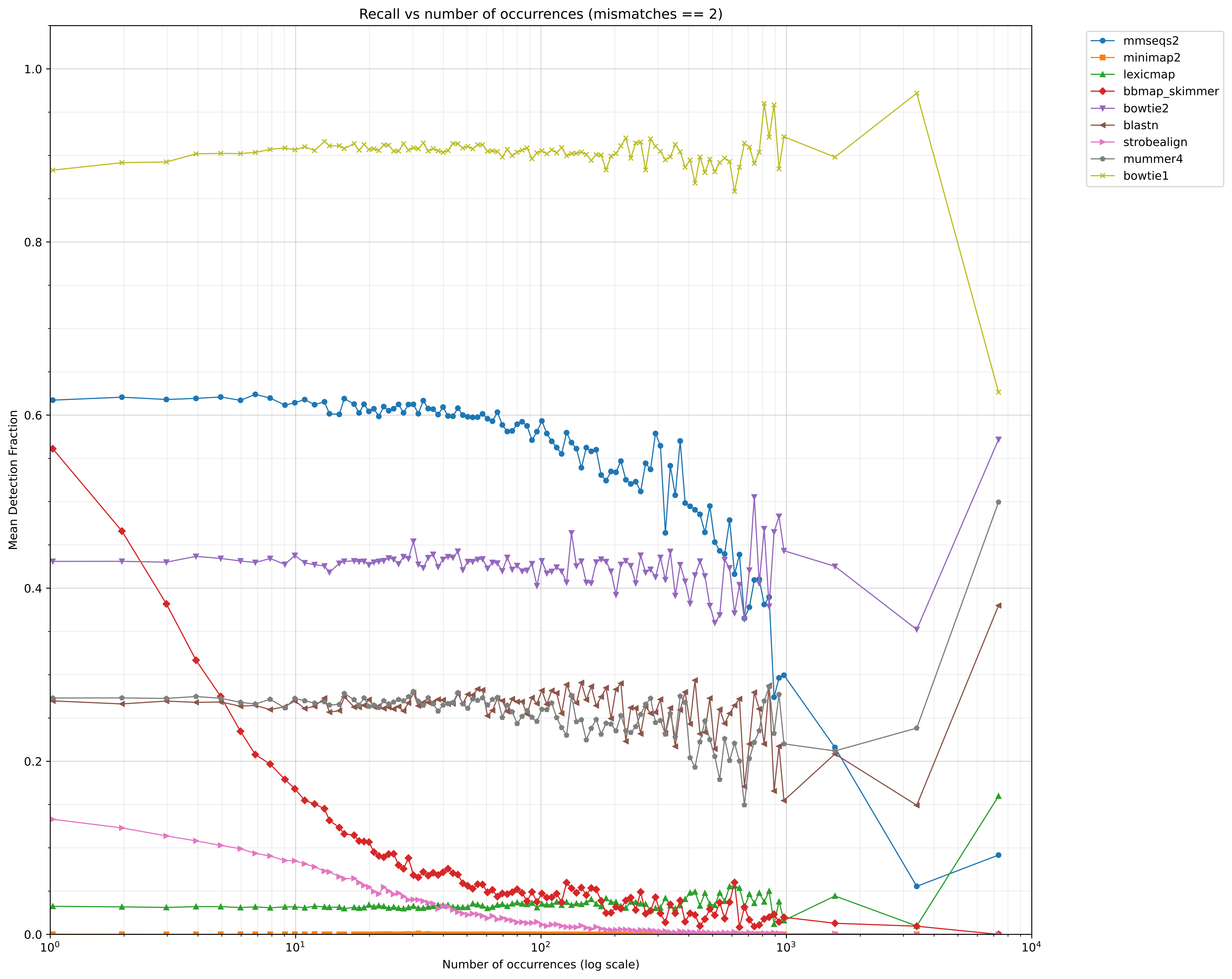

### Supplementary figure 5.D.

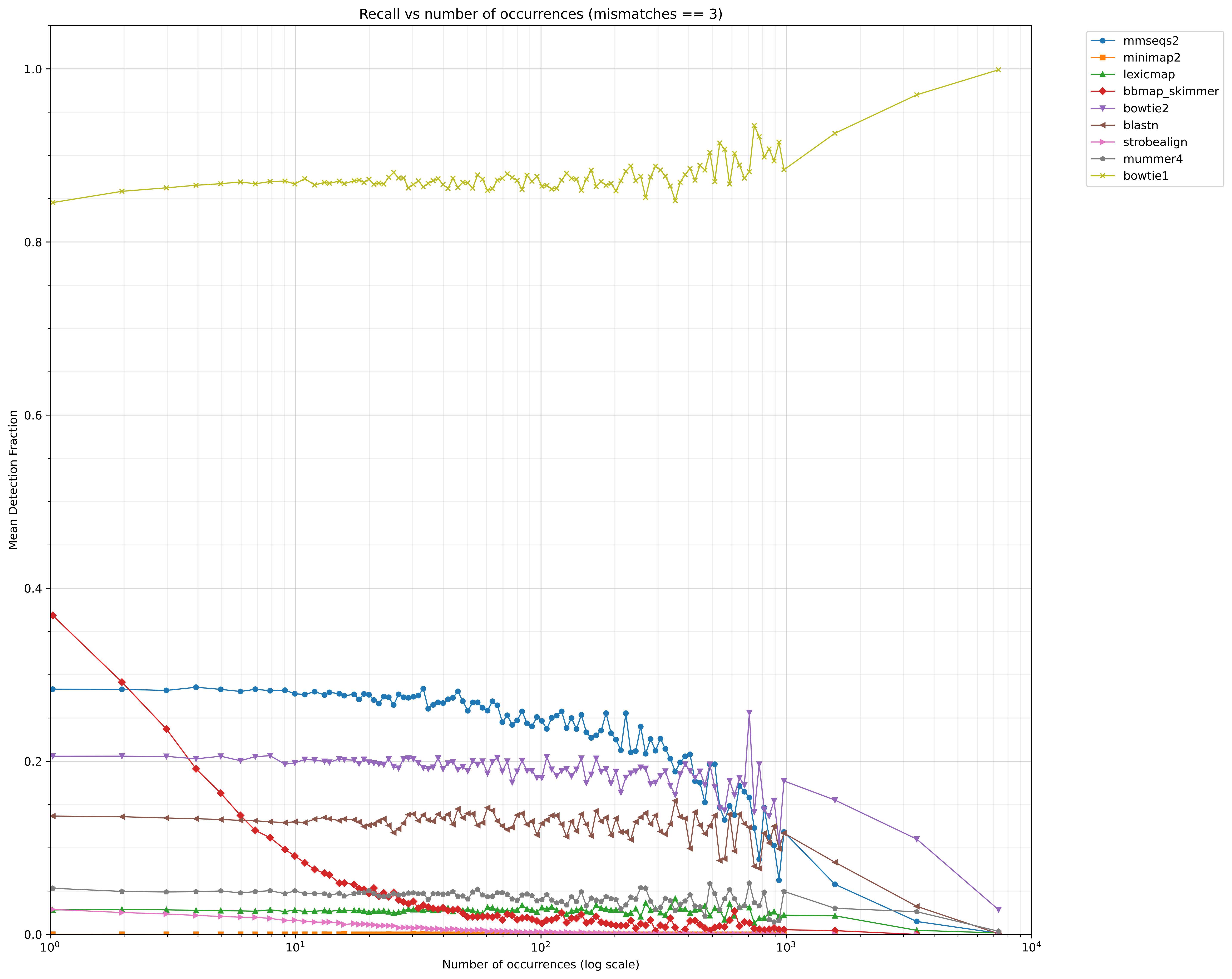

### Supplementary figure 6.1.

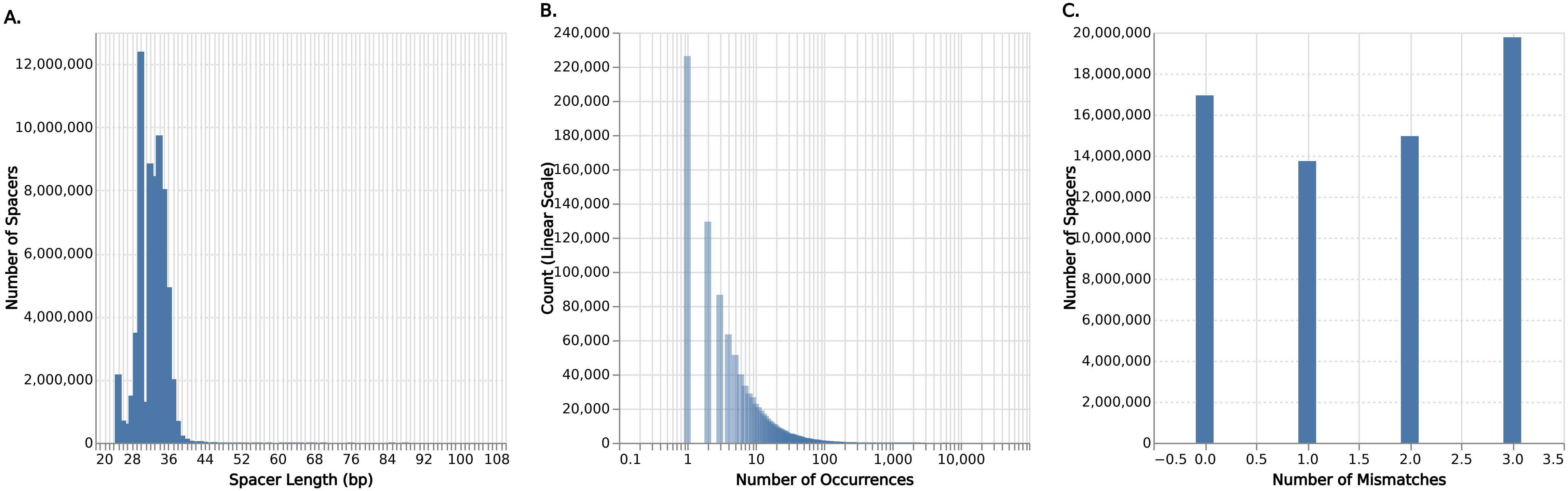

### Supplementary figure 6.2.

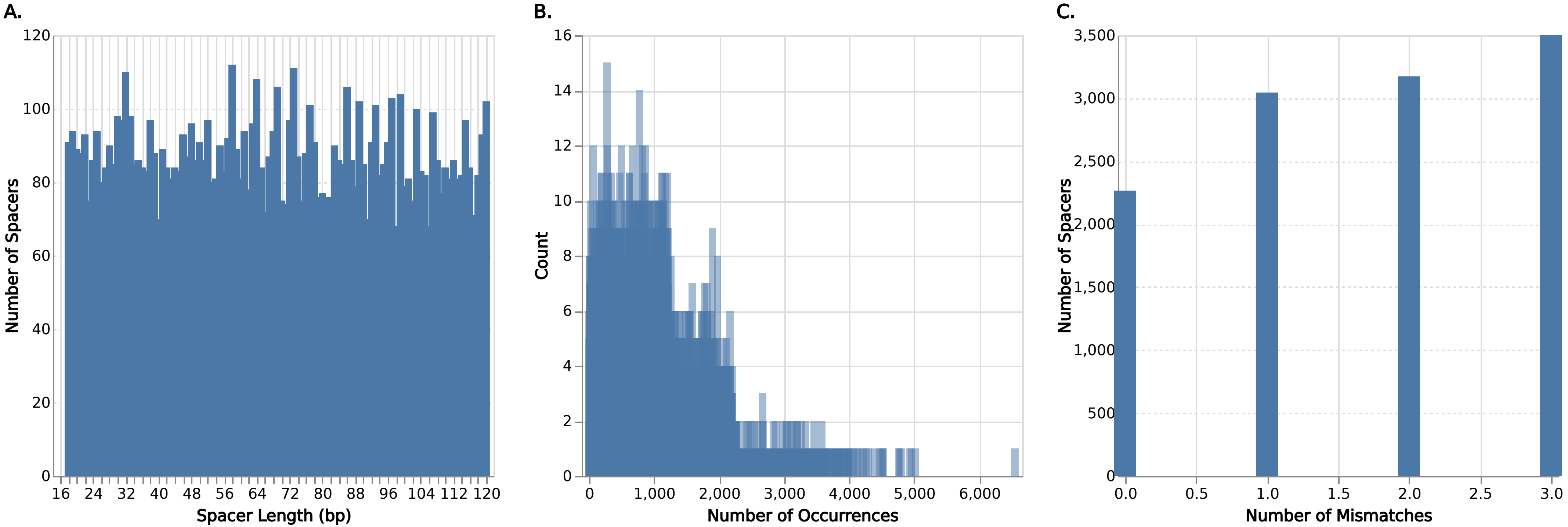

### Supplementary figure 7.A.

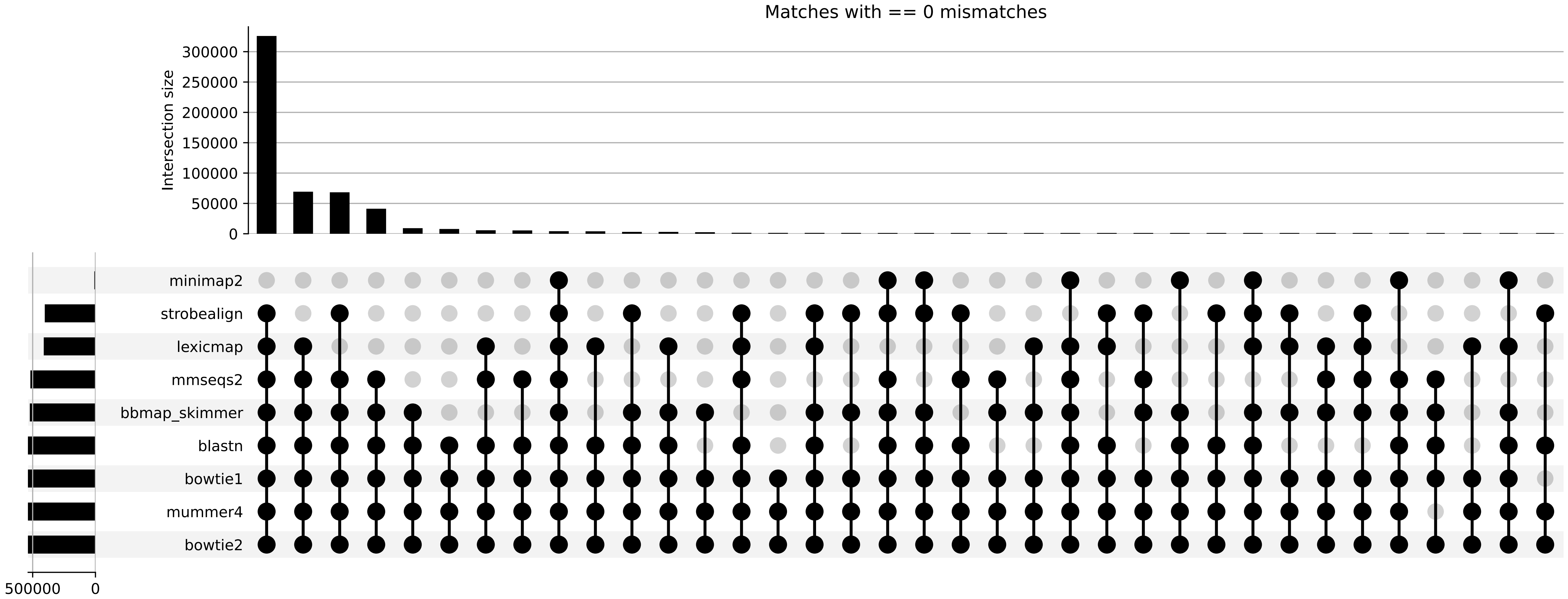

### Supplementary figure 7.B.

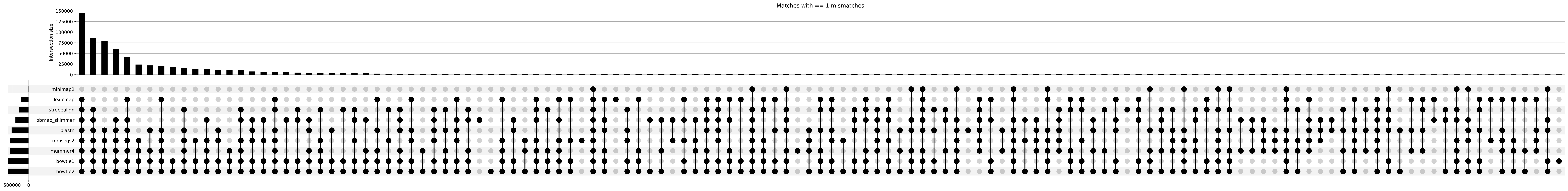

### Supplementary figure 7.C.

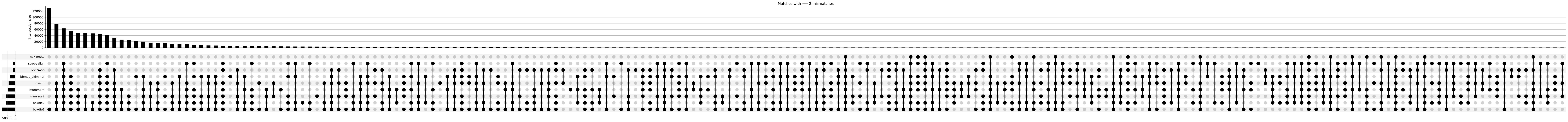

### Supplementary figure 7.D.

#### Matches with == 3 mismatches

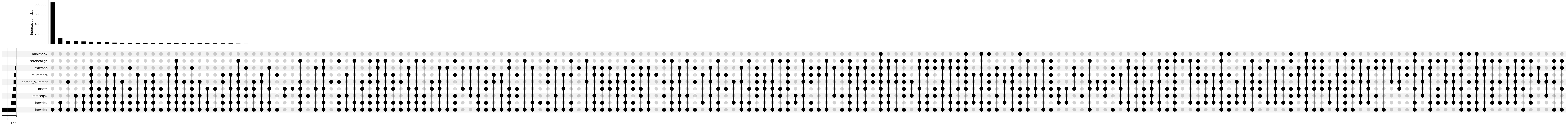
