## Supplementary text for "Tool Choice drastically Impacts CRISPR Spacer-Protospacer Detection"

### Supplementary Materials

#### Tables

##### 1. Tool Configuration Details

For full version and build information see the git repo at [code.jgi.doe.gov/spacersdb/spacer*matching*bench](code.jgi.doe.gov/spacersdb/spacermatchingbench)

| Tool | Version | Command |
|---|---|---|
| bbmapskimmer | 39.13 | `bbmapskimmer.sh sam=1.4 maxindel=0 tipsearch=0 midpad=100 in={spacers_file} ref=./{contigs_file} outm={output_dir}/bbmap_skimmer_output.sam t={threads} minid=0.85 path={output_dir}` |
| blastn | 2.16.0+, Dec 14 2024 | `makeblastdb -in {contigs_file} -dbtype nucl -out {output_dir}/contigs_blastdb``blastn -query {spacers_file} -db {output_dir}/contigs_blastdb -max_target_seqs 1000000 -out {output_dir}/blastn_output.tsv -evalue 1e-5 -num_threads {threads} -task blastn-short -outfmt 6 qaccver saccver nident length mismatch qlen gapopen qstart qend sstart send evalue bitscore""` |
| bowtie1 | bowtie-align-s version 1.3.1 | `bowtie-build --threads {threads} {contigs_file} {results_dir}/simulated_data/bt1_contigs_index``bowtie --threads {threads} -f --all -v 3 -x {results_dir}/simulated_data/bt1_contigs_index {spacers_file} -S {output_dir}/bowtie1_output.sam` |
| bowtie2 | bowtie2-align-s 2.5.4 | `bowtie2-build --large-index --threads {threads} {contigs_file} {results_dir}/simulated_data/contigs_bt2indx``bowtie2 --all --xeq --very-sensitive -x {results_dir}/simulated_data/contigs_bt2indx -f {spacers_file} -S {output_dir}/bowtie2_output.sam --threads {threads}` |
| hisat2 | hisat2-align-s 2.2.1 | `hisat2-build -p {threads} {contigs_file} {output_dir}/hisat2_idx``hisat2 -a --no-spliced-alignment --no-unal --no-softclip --secondary -p {threads} -x {output_dir}/hisat2_idx -f {spacers_file} -S {output_dir}/hisat2_output.sam` |
| lexicmap | v0.5.0 (06741c8) | `mkdir -p {output_dir}/tmp_lexicmap_contigs {output_dir}/tmp_lexicmap_spacers``cp {contigs_file} {output_dir}/tmp_lexicmap_contigs/simulated_contigs.fa``cp {spacers_file} {output_dir}/tmp_lexicmap_spacers/simulated_spacers.fa``lexicmap index -k 15 -m 40000 --seed-max-desert 200 --seed-in-desert-dist 50 -I {output_dir}/tmp_lexicmap_contigs -O {output_dir}/tmp_lexicmap.lmi``lexicmap search -d {output_dir}/tmp_lexicmap.lmi {output_dir}/tmp_lexicmap_spacers/simulated_spacers.fa -o {output_dir}/lexicmap_output.tsv --threads {threads} --align-min-match-len 17 --align-min-match-pident 85 --seed-min-prefix 15 --seed-min-single-prefix 15 --seed-max-dist 100 --seed-max-gap 100 --align-max-gap 20 --align-band 100 --top-n-genomes 0 -a` |
| minimap2 | 2.28-r1209 | `minimap2 -N 100 --eqx -t {threads} -a {contigs_file} {spacers_file} -o {output_dir}/minimap2_output.sam` |
| mmseqs2 | db8ad2d14d0a285ce0ad62bbefd0dce927663315 | `mkdir -p {output_dir}/tmp_spacers {output_dir}/tmp_contigs {output_dir}/tmp_mmseqs {output_dir}/tmp_mmseqs_outputs``mmseqs createdb {spacers_file} {output_dir}/tmp_spacers/mmdb``mmseqs createdb {contigs_file} {output_dir}/tmp_contigs/mmdb``mmseqs search {output_dir}/tmp_spacers/mmdb {output_dir}/tmp_contigs/mmdb {output_dir}/tmp_mmseqs_outputs/mmseqs_output {output_dir}/tmp_mmseqs --min-seq-id 0.85 --min-aln-len 17 --threads {threads} -a --search-type 3 -v` |

| Tool | Version | Command |
|---|---|---|
| | | 1``mmseqs convertalis {output_dir}/tmp_spacers/mmdb {output_dir}/tmp_contigs/mmdb {output_dir}/tmp_mmseqs_outputs/mmseqs_output {output_dir}/mmseqs_output.tsv --format-mode 0 --search-type 4 --format-output query,target,nident,alnlen,mismatch,qlen,gapopen,qstart,qend,tstart,tend,evalue,bits |
| mummer4 | 4.1.0-r1304 // 4.1 | `nucmer --maxmatch --nosimplify --batch=10000000 --threads {threads} --sam-long={output_dir}/mummer4_output.sam -c 1 {contigs_file} {spacers_file}` |
| spacer_containment | 0.1.0 | `spacer-containment --n-threads {threads} {contigs_file} {spacers_file} > {output_dir}/spacer_containment_output.tsv` |
| strobealign | 0.15.0 | `strobealign --eqx -k 15 -N 1000 -t {threads} {contigs_file} {spacers_file} -o {output_dir}/strobealign_output.sam` |

### 2. Recall values for each tool at different mismatch thresholds

#### A. IMG/VR4 dataset

Every row lists the results for a specific mismatch threshold and tool.
The values are for exact mismatches (not max), and represent the total number of unique spacer-contig pairs (aligning at that mismatch threshold).
The fraction is tool*matches divided by total*possible.

| mismatches | tool | total_possible | tool_matches | fraction |
|---|---|---|---|---|
| 0 | mummer4 | 16866829 | 16866546 | 0.9999832 |
| 0 | mmseqs2 | 16866829 | 15803286 | 0.9369447 |
| 0 | bowtie1 | 16866829 | 16866784 | 0.9999973 |
| 0 | lexicmap | 16866829 | 6033115 | 0.3576911 |
| 0 | minimap2 | 16866829 | 5110 | 0.000303 |
| 0 | strobealign | 16866829 | 4233236 | 0.25098 |
| 0 | blastn | 16866829 | 16850968 | 0.9990596 |
| 0 | bbmap_skimmer | 16866829 | 14978104 | 0.8880213 |
| 0 | bowtie2 | 16866829 | 16866667 | 0.9999904 |
| 1 | mummer4 | 12197007 | 8032640 | 0.6585747 |
| 1 | mmseqs2 | 12197007 | 9805301 | 0.8039104 |
| 1 | bowtie1 | 12197007 | 11992384 | 0.9832235 |
| 1 | lexicmap | 12197007 | 835330 | 0.0684865 |
| 1 | minimap2 | 12197007 | 1086 | 0.000089038237 |
| 1 | strobealign | 12197007 | 1311189 | 0.1075009 |
| 1 | blastn | 12197007 | 7318398 | 0.6000159 |
| 1 | bbmap_skimmer | 12197007 | 2413864 | 0.1979063 |
| 1 | bowtie2 | 12197007 | 12112313 | 0.9930562 |
| 2 | mummer4 | 12359867 | 3129926 | 0.253233 |
| 2 | mmseqs2 | 12359867 | 6756086 | 0.5466148 |
| 2 | bowtie1 | 12359867 | 11173197 | 0.9039901 |
| 2 | lexicmap | 12359867 | 426554 | 0.0345112 |
| 2 | minimap2 | 12359867 | 667 | 0.000053964982 |
| 2 | strobealign | 12359867 | 375251 | 0.0303604 |
| 2 | blastn | 12359867 | 3217012 | 0.2602789 |
| 2 | bbmap_skimmer | 12359867 | 1047205 | 0.0847262 |
| 2 | bowtie2 | 12359867 | 5243454 | 0.4242322 |
| 3 | mummer4 | 16472212 | 673444 | 0.0408836 |
| 3 | mmseqs2 | 16472212 | 3560155 | 0.216131 |
| 3 | bowtie1 | 16472212 | 14523177 | 0.8816774 |
| 3 | lexicmap | 16472212 | 430208 | 0.0261172 |
| 3 | minimap2 | 16472212 | 392 | 0.000023797654 |

| mismatches | tool | total_possible | tool_matches | fraction |
|---|---|---|---|---|
| 3 | strobealign | 16472212 | 88246 | 0.0053573 |
| 3 | blastn | 16472212 | 1972148 | 0.1197258 |
| 3 | bbmap_skimmer | 16472212 | 630914 | 0.0383017 |
| 3 | bowtie2 | 16472212 | 2998722 | 0.1820473 |

### B. Synthetic dataset

| mismatches | tool | total_possible | tool_matches | recall | false_positives | false_negatives | precision | f1_score |
|---|---|---|---|---|---|---|---|---|
| 0 | minimap2 | 1899024 | 176 | 0.00009267918678 | 49 | 1898848 | 0.7822222222 | 0.0001853364146 |
| 0 | bbmapskimmer | 1899024 | 377426 | 0.1987473565 | 1010 | 1521598 | 0.9973311207 | 0.3314446796 |
| 0 | mmseqs2map | 1899024 | 1021533 | 0.5379252711 | 57809 | 877491 | 0.9464405165 | 0.6859687493 |
| 0 | mmseqs2 | 1899024 | 1021533 | 0.5379252711 | 134608 | 877491 | 0.8835712945 | 0.6687252571 |
| 0 | minimap2_mod | 1899024 | 176 | 0.00009267918678 | 49 | 1898848 | 0.7822222222 | 0.0001853364146 |
| 0 | bowtie1 | 1899024 | 1899024 | 1 | 0 | 0 | 1 | 1 |
| 0 | bbmapskimmermod | 1899024 | 1899024 | 1 | 1130 | 0 | 0.9994053114 | 0.9997025672 |
| 0 | bwa_mem | 1899024 | 683227 | 0.3597779702 | 15626 | 1215797 | 0.9776405052 | 0.5259887208 |
| 0 | lexicmap | 1899024 | 1427612 | 0.7517609045 | 469844 | 471412 | 0.7523821369 | 0.7520713924 |
| 0 | hisat2 | 1899024 | 1875654 | 0.9876936784 | 3 | 23370 | 0.9999984006 | 0.9938079536 |
| 0 | minimap2_og | 1899024 | 48 | 0.00002527614185 | 6 | 1898976 | 0.8888888889 | 0.00005055084625 |
| 0 | spacer_containment | 1899024 | 1899024 | 1 | 0 | 0 | 1 | 1 |
| 0 | bowtie2 | 1899024 | 1899024 | 1 | 3 | 0 | 0.9999984202 | 0.9999992101 |
| 0 | blastn | 1899024 | 1745238 | 0.919018401 | 596453 | 153786 | 0.7452896219 | 0.8230866729 |
| 0 | mummer4 | 1899024 | 1860393 | 0.9796574451 | 250245 | 38631 | 0.881436324 | 0.9279550246 |
| 0 | strobealign | 1899024 | 42016 | 0.0221250495 | 193 | 1857008 | 0.9954275155 | 0.04328795152 |
| 1 | minimap2 | 1992339 | 86 | 0.00004316534485 | 49 | 1992253 | 0.637037037 | 0.00008632484037 |
| 1 | bbmapskimmer | 1992339 | 814246 | 0.4086884812 | 1010 | 1178093 | 0.9987611253 | 0.5800309518 |
| 1 | mmseqs2map | 1992339 | 982267 | 0.4930220209 | 57809 | 1010072 | 0.9444184848 | 0.6478447046 |
| 1 | mmseqs2 | 1992339 | 971696 | 0.4877161969 | 134608 | 1020643 | 0.8783263913 | 0.627175186 |
| 1 | minimap2_mod | 1992339 | 86 | 0.00004316534485 | 49 | 1992253 | 0.637037037 | 0.00008632484037 |
| 1 | bowtie1 | 1992339 | 1924529 | 0.9659646275 | 0 | 67810 | 1 | 0.9826876984 |
| 1 | bbmapskimmermod | 1992339 | 1903036 | 0.9551768048 | 1130 | 89303 | 0.9994065643 | 0.9767912527 |
| 1 | bwa_mem | 1992339 | 569097 | 0.2856426542 | 15626 | 1423242 | 0.9732762351 | 0.4416634136 |
| 1 | lexicmap | 1992339 | 848885 | 0.4260745787 | 469844 | 1143454 | 0.643714516 | 0.512756005 |
| 1 | hisat2 | 1992339 | 1638156 | 0.8222275426 | 3 | 354183 | 0.9999981687 | 0.9024414832 |
| 1 | minimap2_og | 1992339 | 12 | 0.00000602 | 6 | 1992327 | 0.6666666667 | 0.00001204603392 |
| 1 | spacer_containment | 0 | 0 | 0 | 0 | 0 | 0 | 0 |
| 1 | bowtie2 | 1992339 | 1924529 | 0.9659646275 | 3 | 67810 | 0.9999984412 | 0.9826869458 |
| 1 | blastn | 1992339 | 1534121 | 0.7700100234 | 596453 | 458218 | 0.7200505591 | 0.7441927589 |
| 1 | mummer4 | 1992339 | 1605268 | 0.8057203117 | 250245 | 387071 | 0.8651343321 | 0.8343709685 |
| 1 | strobealign | 1992339 | 39698 | 0.01992532395 | 193 | 1952641 | 0.9951618159 | 0.03906841253 |
| 2 | minimap2 | 2137067 | 91 | 0.00004258172533 | 49 | 2136976 | 0.65 | 0.00008515787193 |
| 2 | bbmapskimmer | 2137067 | 804812 | 0.3765965222 | 1010 | 1332255 | 0.9987466215 | 0.5469536907 |
| 2 | mmseqs2map | 2137067 | 912682 | 0.4270722443 | 57809 | 1224385 | 0.9404332446 | 0.5873949899 |
| 2 | mmseqs2 | 2137067 | 901193 | 0.4216961845 | 134608 | 1235874 | 0.8700445356 | 0.5680620814 |
| 2 | minimap2_mod | 2137067 | 91 | 0.00004258172533 | 49 | 2136976 | 0.65 | 0.00008515787193 |
| 2 | bowtie1 | 2137067 | 1934586 | 0.9052528536 | 0 | 202481 | 1 | 0.9502705658 |
| 2 |  | 2137067 | 1734297 | 0.811531412 | 1130 | 402770 | 0.9993488634 | 0.8957002903 |

| mismatches | tool | total_possible | tool_matches | recall | false_positives | false_negatives | precision | f1_score |
|---|---|---|---|---|---|---|---|---|
| | bbmapskimmermod | | | | | | | |
| 2 | bwa_mem | 2137067 | 515856 | 0.2413850385 | 15626 | 1621211 | 0.9705991924 | 0.3866190952 |
| 2 | lexicmap | 2137067 | 562385 | 0.2631574022 | 469844 | 1574682 | 0.544825809 | 0.3548958507 |
| 2 | hisat2 | 2137067 | 1115625 | 0.5220355749 | 3 | 1021442 | 0.9999973109 | 0.6859696344 |
| 2 | minimap2_og | 2137067 | 26 | 0.00001216620724 | 6 | 2137041 | 0.8125 | 0.00002433205013 |
| 2 | spacer_containment | 0 | 0 | 0 | 0 | 0 | 0 | 0 |
| 2 | bowtie2 | 2137067 | 1573821 | 0.7364397092 | 3 | 563246 | 0.9999980938 | 0.8482173149 |
| 2 | blastn | 2137067 | 1355458 | 0.6342608819 | 596453 | 781609 | 0.6944261291 | 0.6629813122 |
| 2 | mummer4 | 2137067 | 1432751 | 0.6704286763 | 250245 | 704316 | 0.851309807 | 0.7501190425 |
| 2 | strobealign | 2137067 | 34729 | 0.01625077735 | 193 | 2102338 | 0.9944733979 | 0.03197898332 |
| 3 | minimap2 | 2609997 | 220 | 0.00008429128463 | 49 | 2609777 | 0.8178438662 | 0.000168565196 |
| 3 | bbmapskimmer | 2609997 | 760618 | 0.291424856 | 1010 | 1849379 | 0.9986738933 | 0.4511877804 |
| 3 | mmseqs2map | 2609997 | 860977 | 0.3298766244 | 57809 | 1749020 | 0.9370811048 | 0.4879738992 |
| 3 | mmseqs2 | 2609997 | 862278 | 0.3303750924 | 134608 | 1747719 | 0.8649715213 | 0.4781291769 |
| 3 | minimap2_mod | 2609997 | 228 | 0.00008735642225 | 49 | 2609769 | 0.8231046931 | 0.0001746943041 |
| 3 | bowtie1 | 2609997 | 2238309 | 0.8575906409 | 0 | 371688 | 1 | 0.9233365221 |
| 3 | bbmapskimmermod | 2609997 | 1534102 | 0.5877792197 | 1130 | 1075895 | 0.9992639549 | 0.7401772013 |
| 3 | bwa_mem | 2609997 | 497712 | 0.1906944721 | 15626 | 2112285 | 0.969560017 | 0.3187054863 |
| 3 | lexicmap | 2609997 | 443019 | 0.1697392756 | 469844 | 2166978 | 0.4853072148 | 0.2515109882 |
| 3 | hisat2 | 2609997 | 564049 | 0.2161109764 | 3 | 2045948 | 0.9999946813 | 0.3554129127 |
| 3 | minimap2_og | 2609997 | 35 | 0.0000134099771 | 6 | 2609962 | 0.8536585366 | 0.0000268195329 |
| 3 | spacer_containment | 0 | 0 | 0 | 0 | 0 | 0 | 0 |
| 3 | bowtie2 | 2609997 | 1415300 | 0.5422611597 | 3 | 1194697 | 0.9999978803 | 0.7032022458 |
| 3 | blastn | 2609997 | 1270806 | 0.4868994102 | 596453 | 1339191 | 0.6805729682 | 0.5676718061 |
| 3 | mummer4 | 2609997 | 1338301 | 0.5127595932 | 250245 | 1271696 | 0.842469151 | 0.6375073448 |
| 3 | strobealign | 2609997 | 30913 | 0.01184407492 | 193 | 2579084 | 0.9937954092 | 0.02340915898 |

### Figures

#### Supplementary figure 1.

Benchmarking framework pipeline overview.

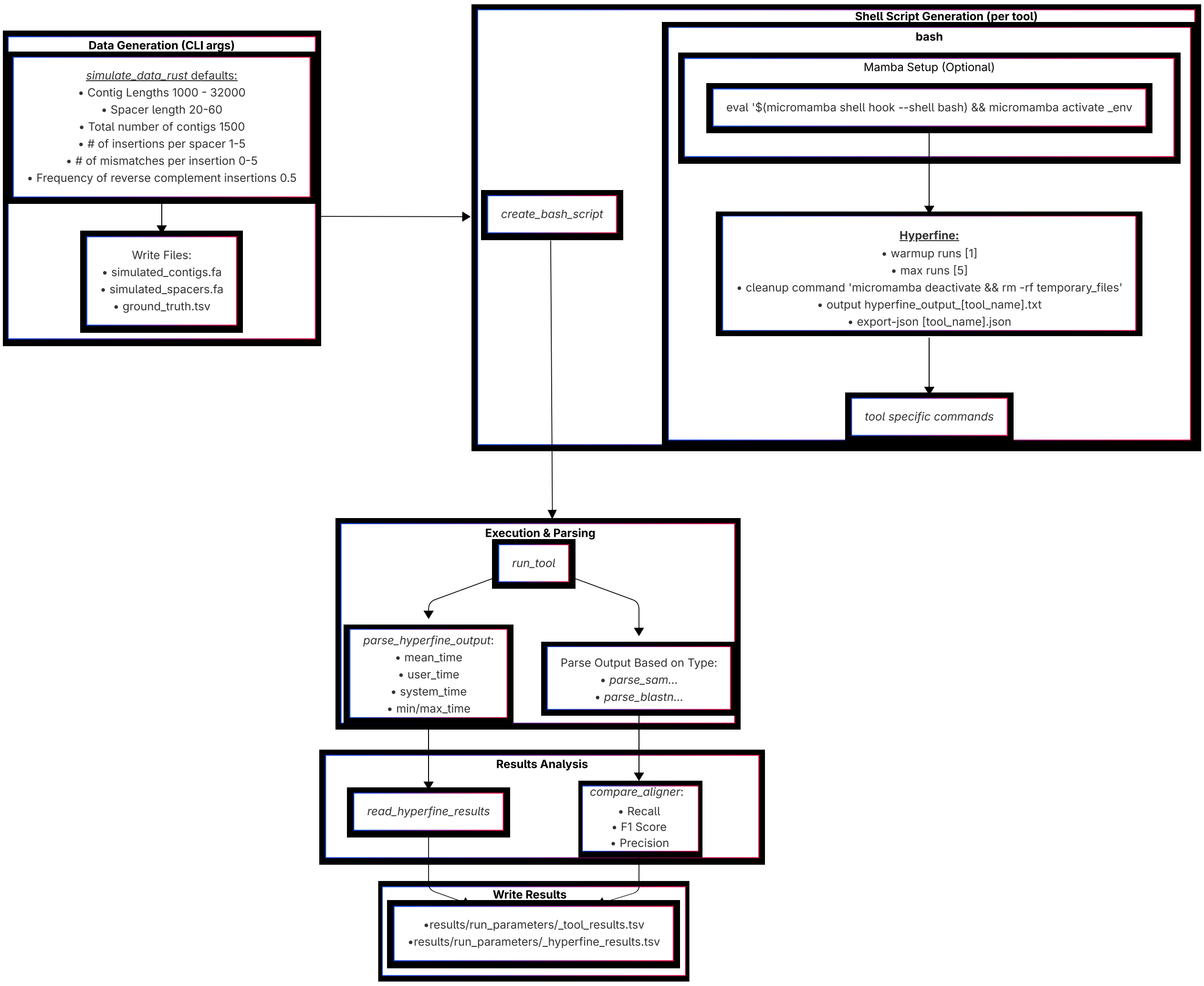


### Supplementary figure 2.

IMG/VR v4 based results in a tool vs tool comparison. Unlike the similar matrix in the main text(figure 1) which showcases the values for up to 1 and 3 mismatches, here the matrixes are separated for each mismatch threshold at an **exact** mismatch value. From top left to bottom right, the mismatch threshold is 0, 1, 2, 3. Like the main text figure, the value of a cell(i,j) is the fraction of spacer-contig pairs identified by the tool listed in row i, which were not identified by the tool listed in the j column.

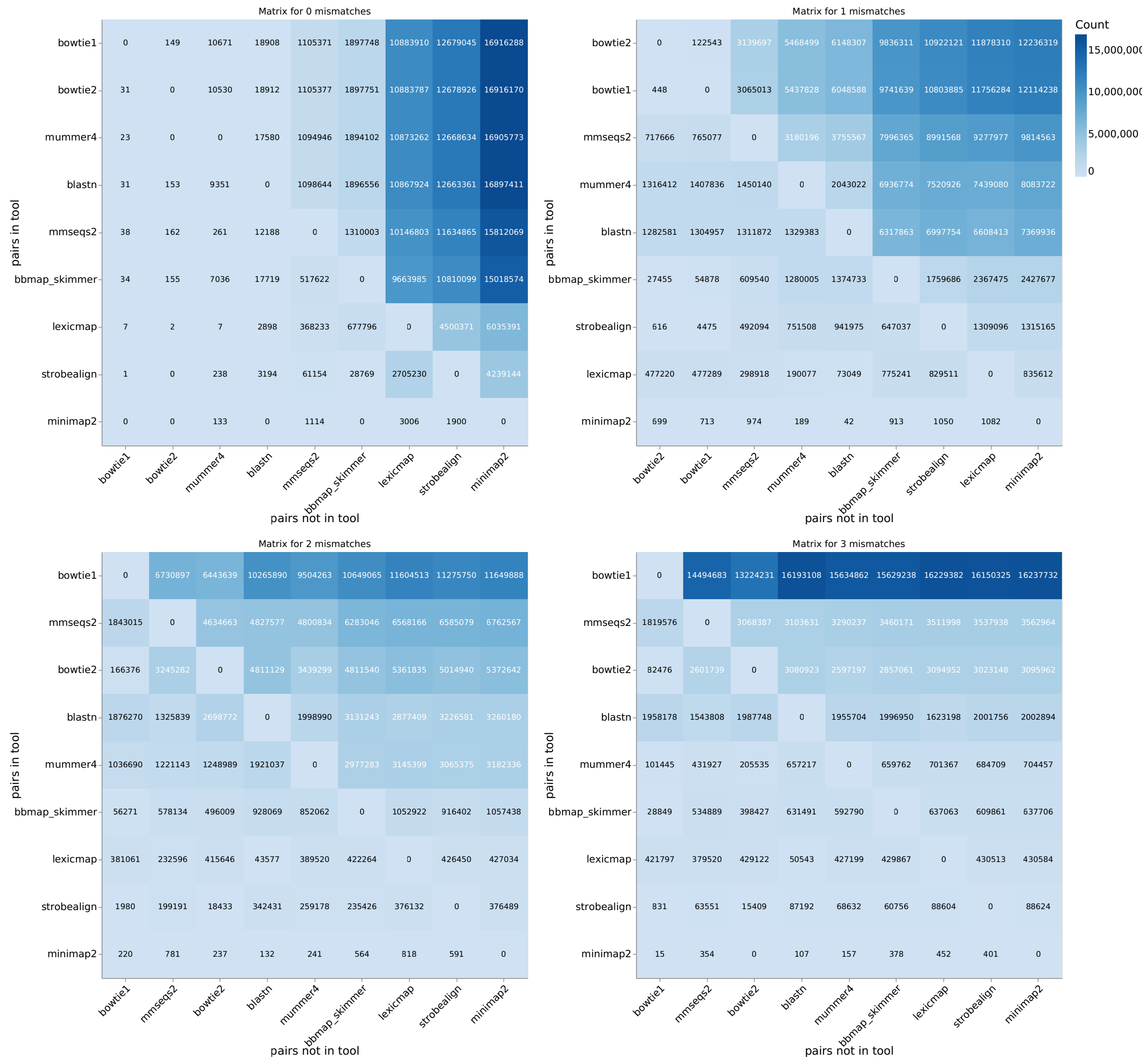


### Supplementary figure 3

Simulated dataset performance metrics.
Percision = True Positives / (True Positives + False Positives)
Recall = True Positives / (True Positives + False Negatives)
F1 = 2 * (Percision * Recall) / (Percision + Recall)
Note: Because of the various prefiltering steps, the number of False Negatives and False Positives may not be indicative of the actual raw tool-reported results. As such, we recommend focusing on the recall rate between tools, which is equivalent to the fraction of detected spacer-protospacer pairs out of all the spacer-protospacer pairs in the reference file.

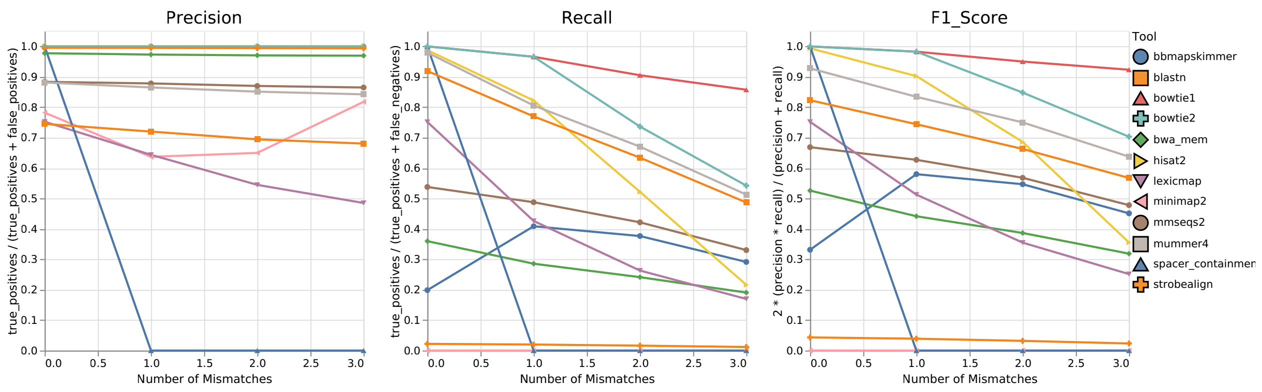


### Supplementary figure 4.

**Simulated dataset** recall (detection fraction) for different values of spacer occurrences.

#### A. for exact mismatches, in a contig dependent manner

Per contigs manner means the that the recall measure is the fraction of occurences each tool identified out of the total number of spacer-contig pairs, identified by all tools.

1. 0 mismatches

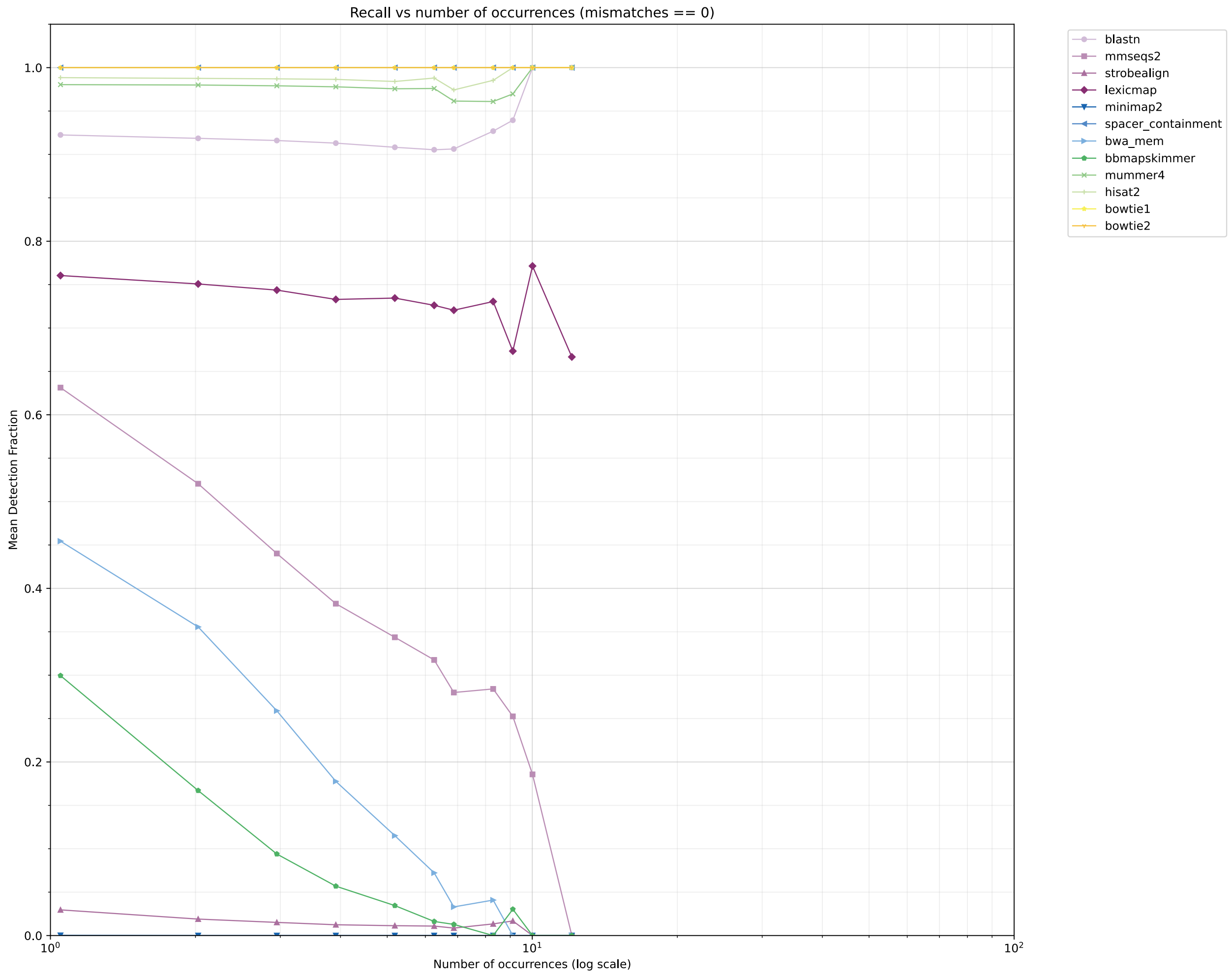


1. exactly 1 mismatch

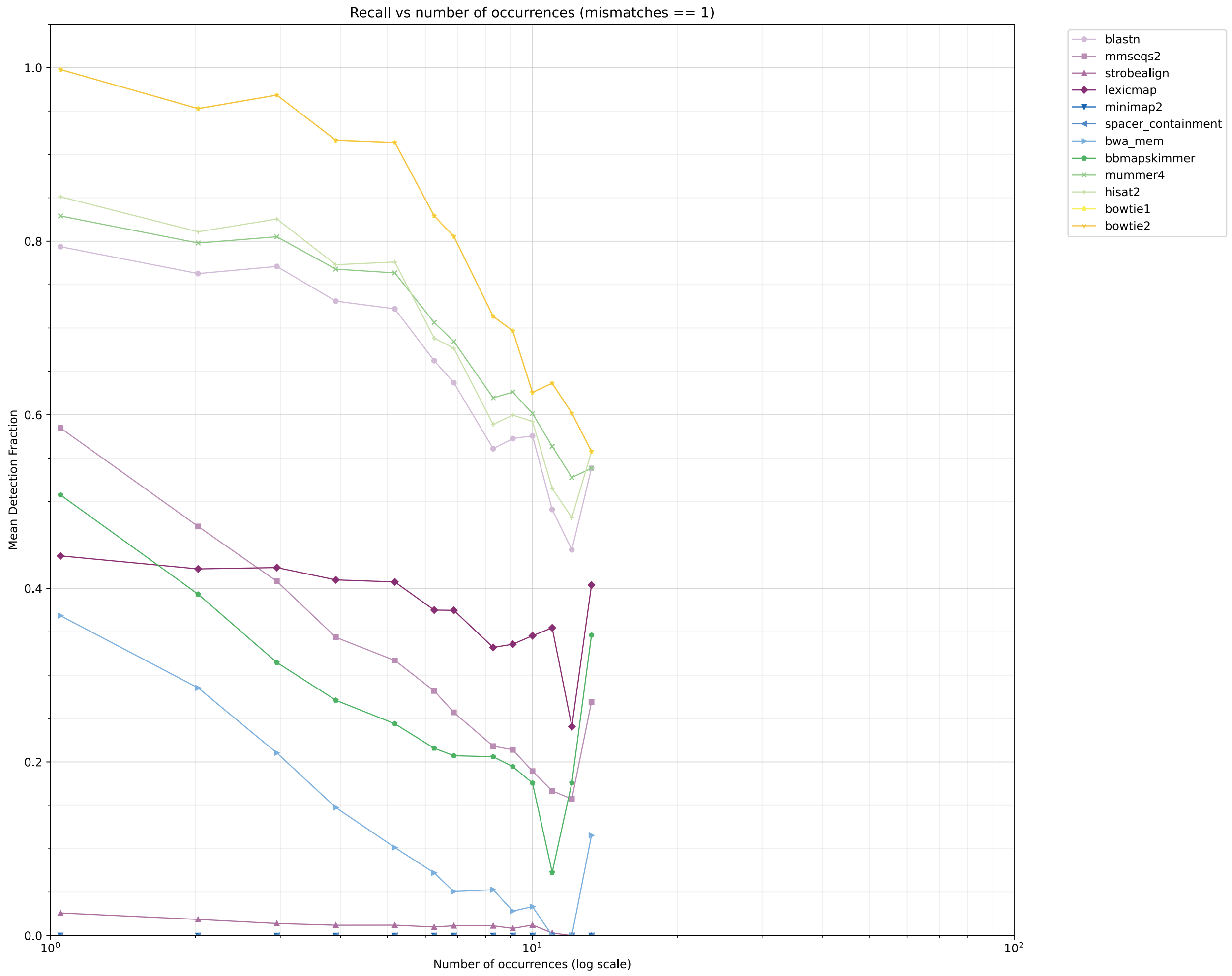


1. exactly 2 mismatches

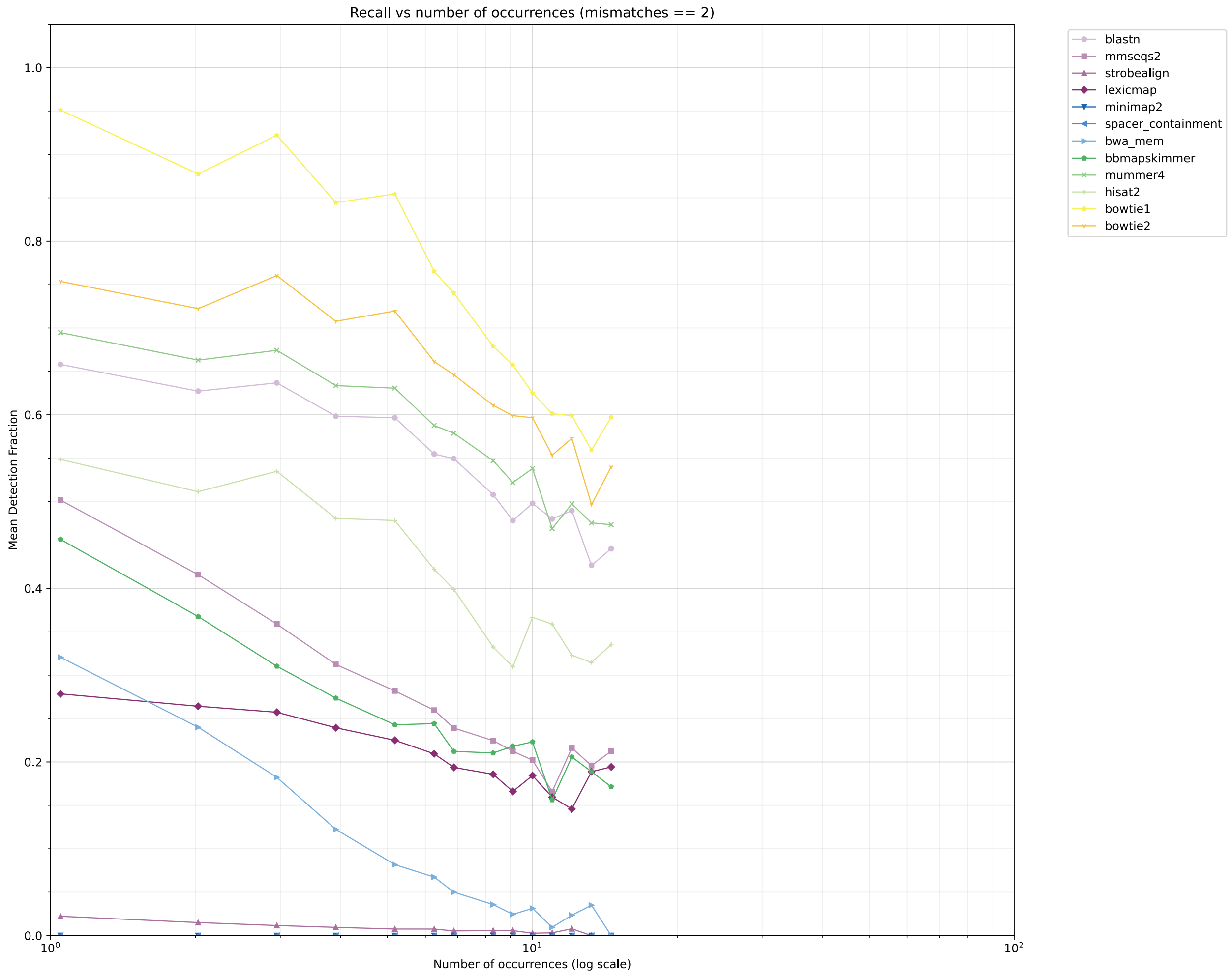


1. exactly 3 mismatches

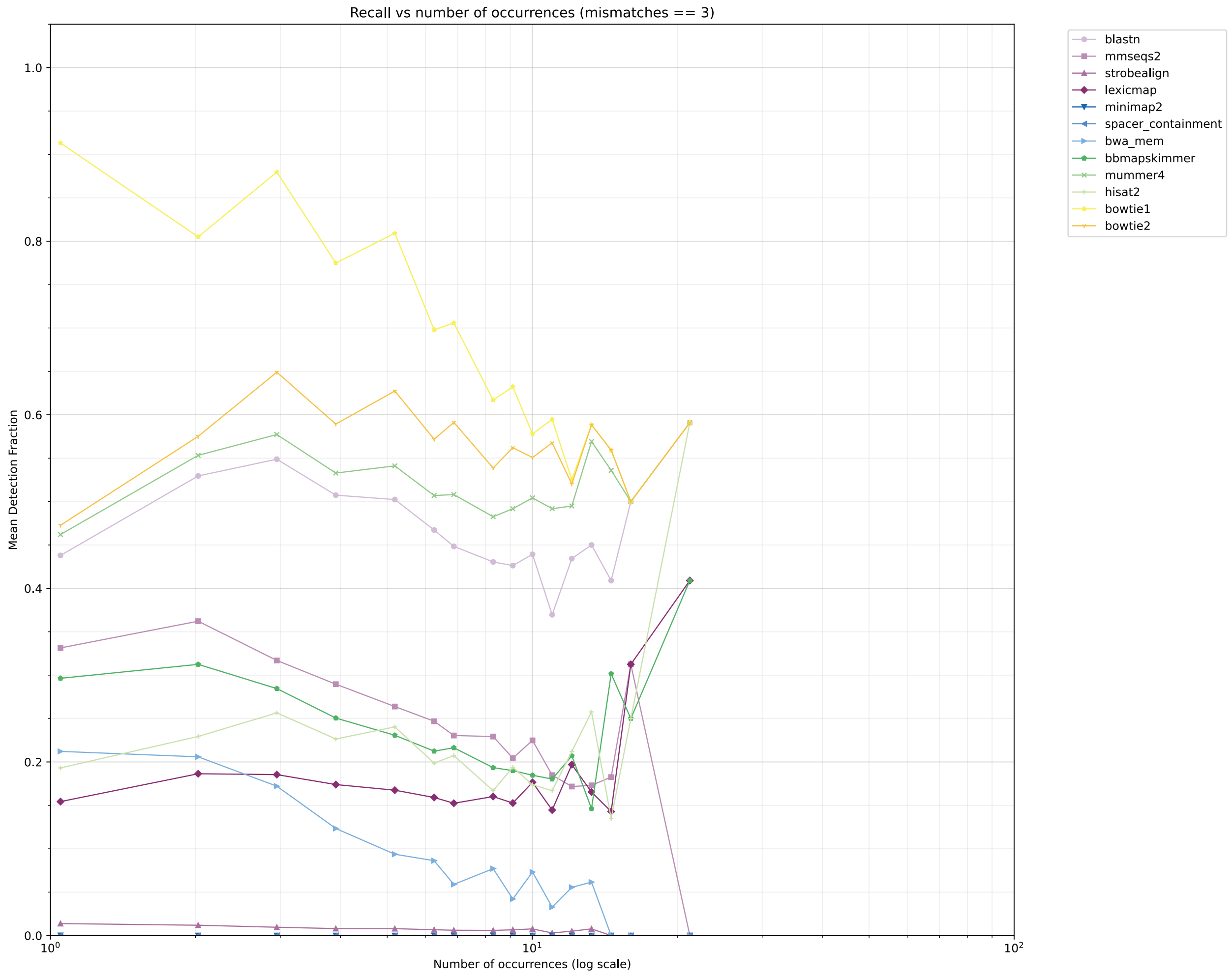


### B. up to 1 and 3 mismatches, in a contig independent manner

Per contig independent manner means that the recall measure is the fraction of occurences each tool identified out of the total number of times the spacer occurs in the reference file (regardless of in which contig).

1. up to 1 mismatches

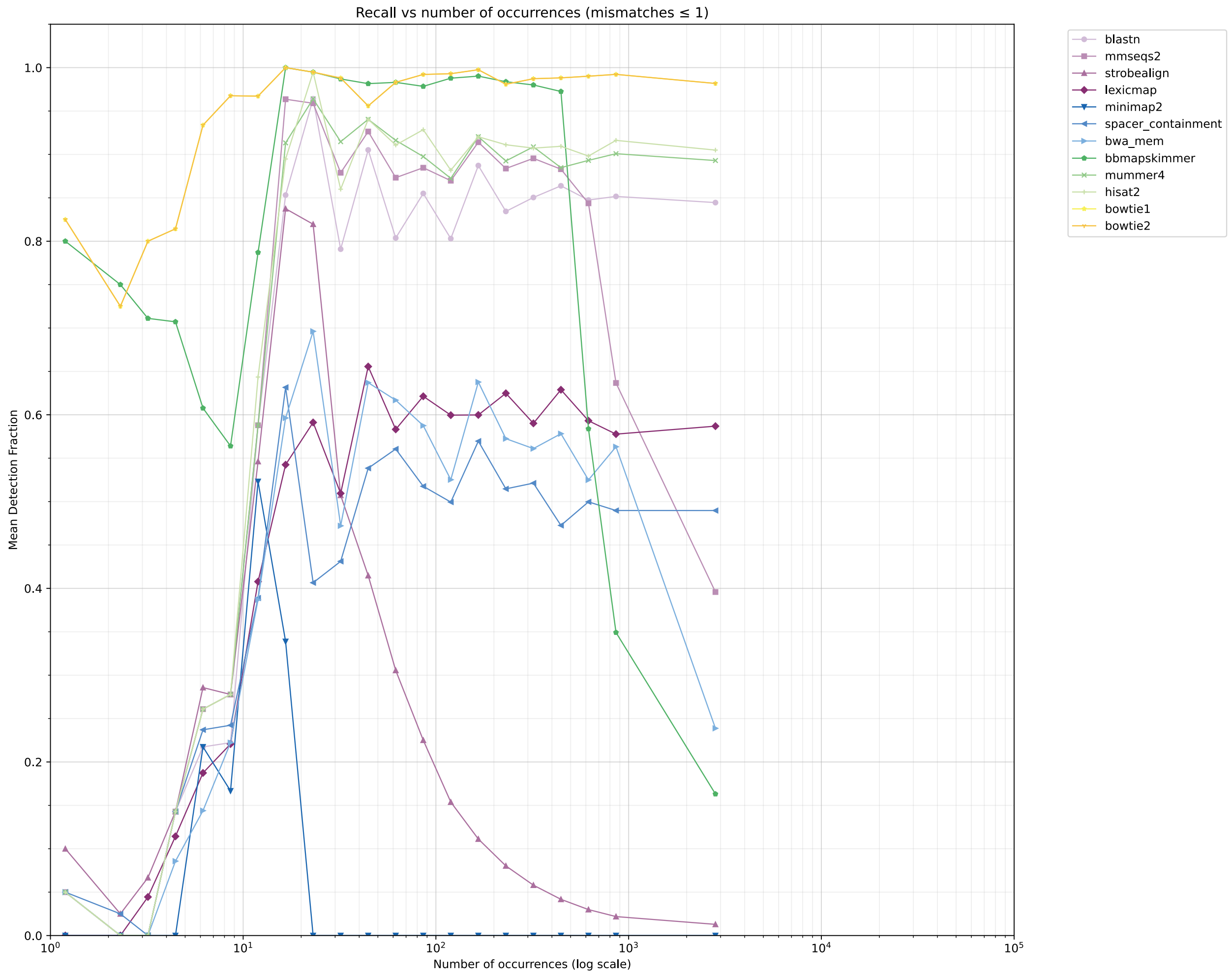


1. up to 3 mismatches

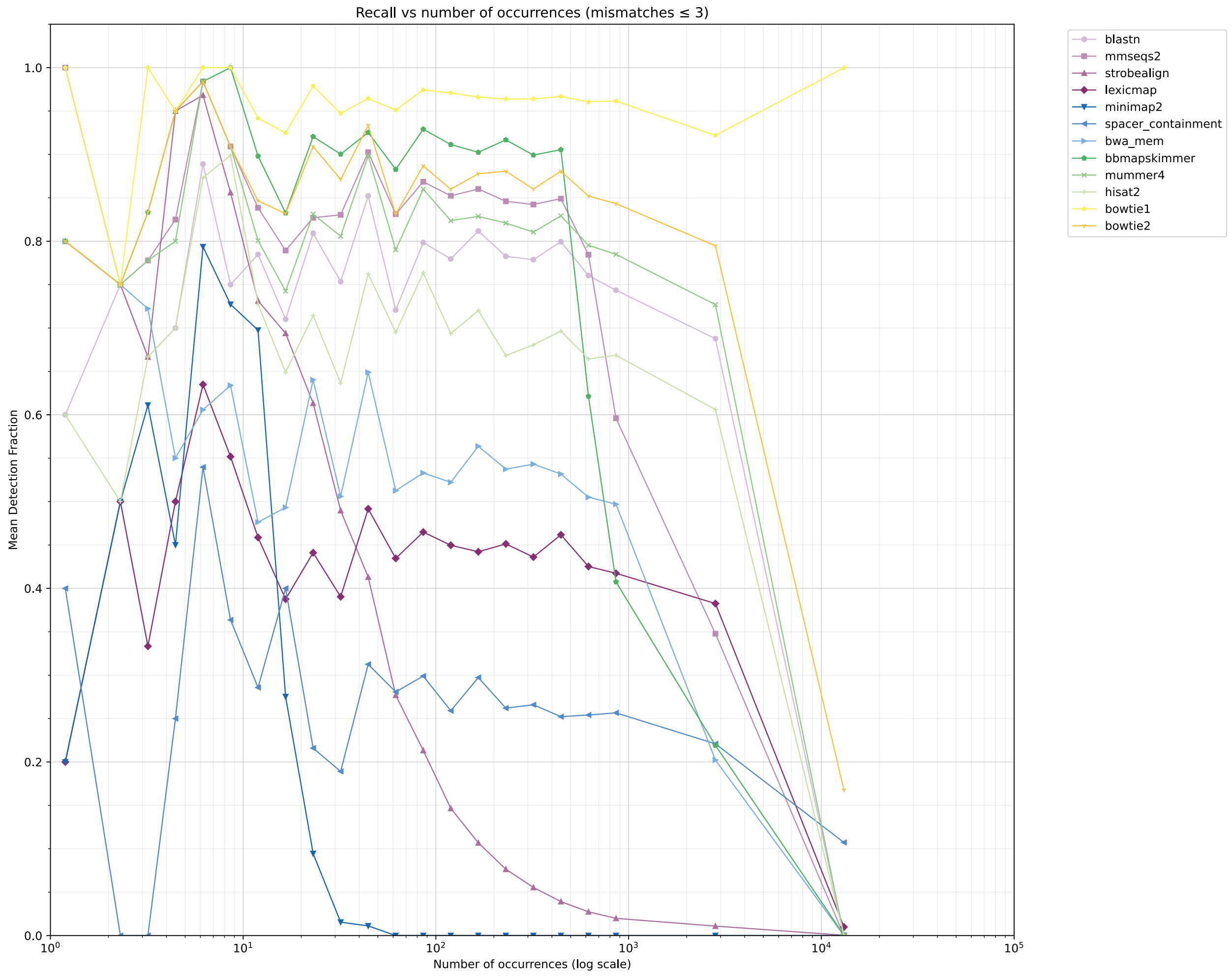


### Supplementary figure 5.

Tool recall versus occurrence frequency for **IMG/VR4 dataset**.
Similar to main text figure, except for exact values for different mismatch thresholds (0-3).

### A. 0 mismatches

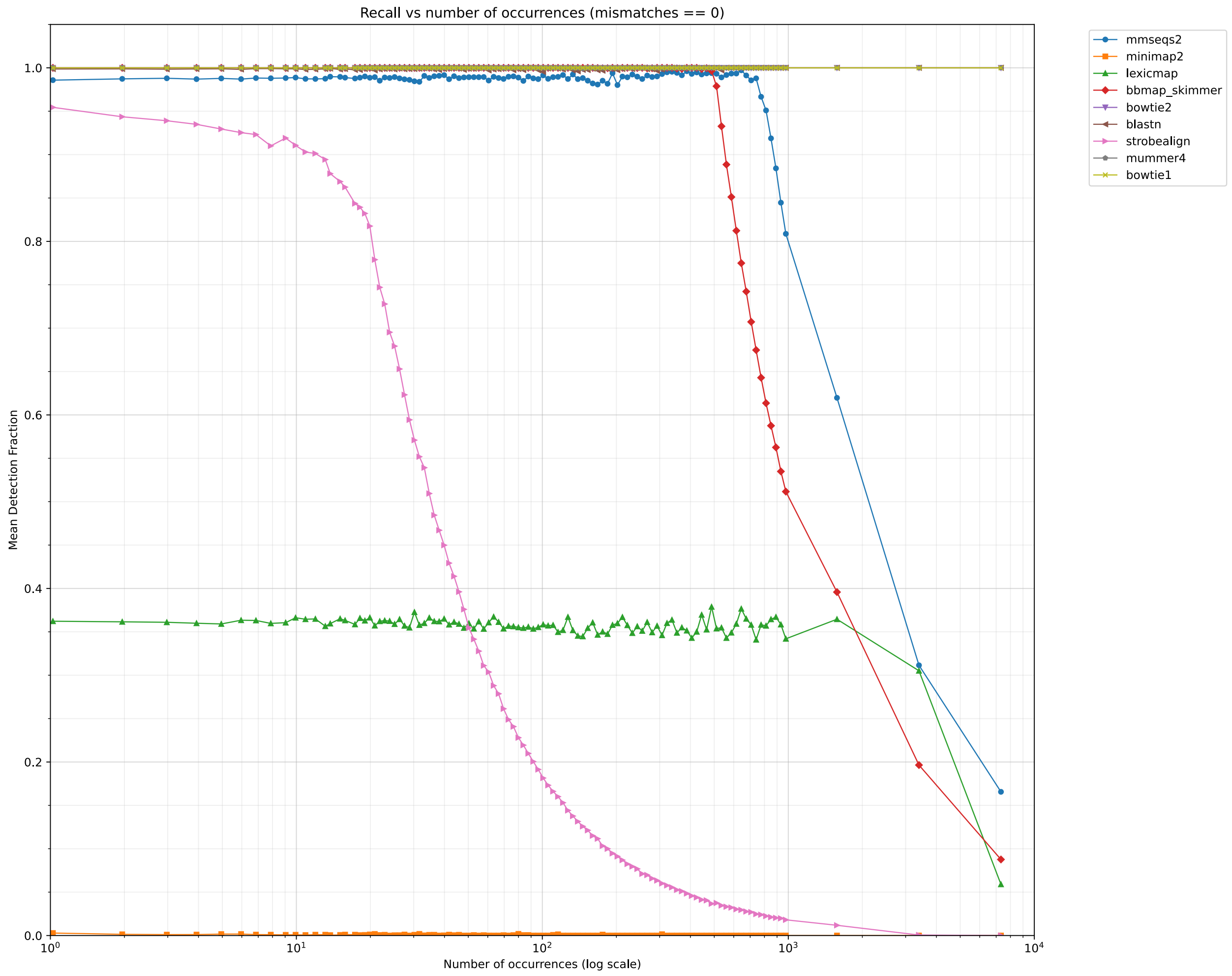

### B. 1 mismatch

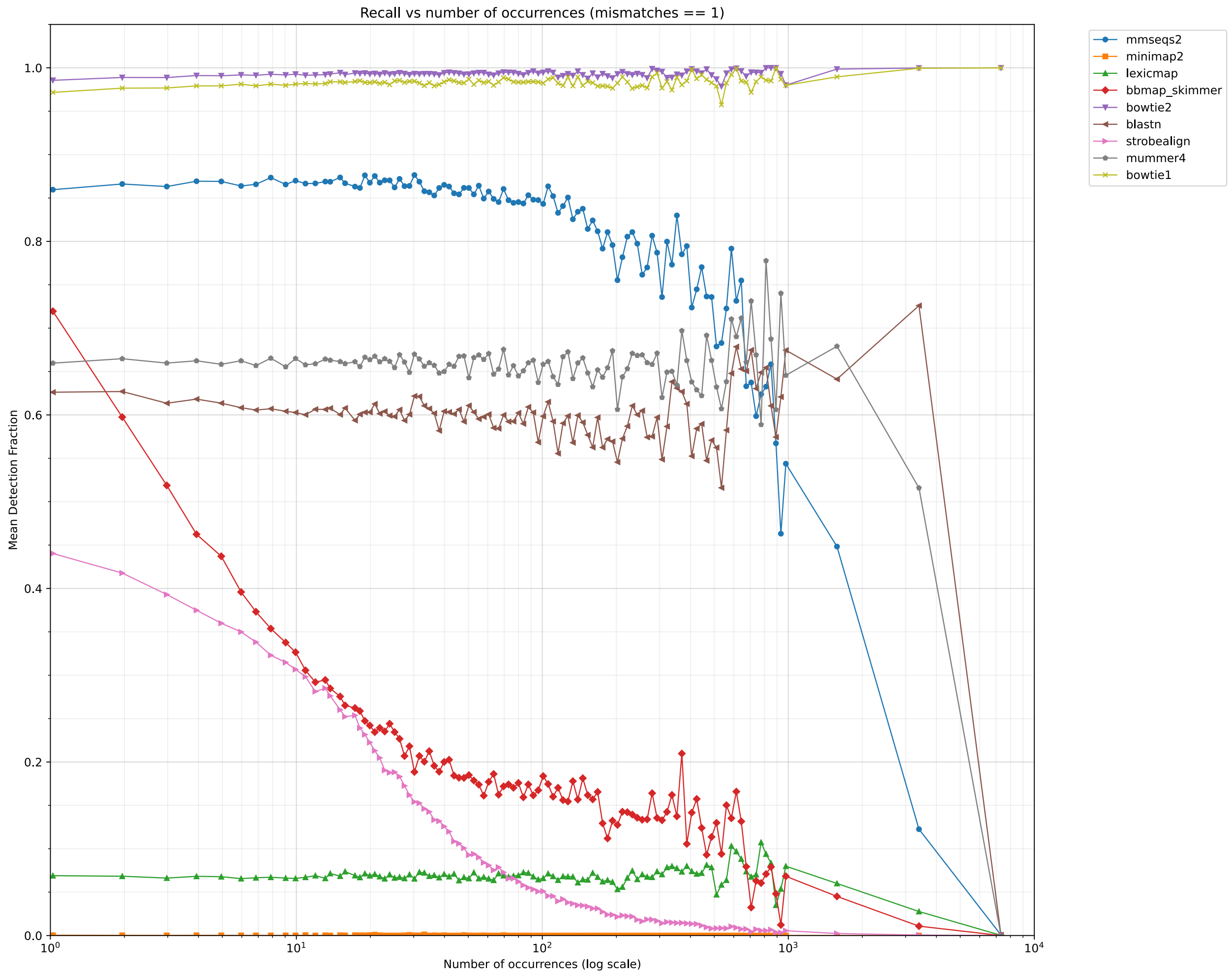

Recall vs number of occurrences (mismatches == 1)
Mean Detection Fraction
Number of occurrences (log scale)
0.0
0.2
0.4
0.6
0.8
1.0
$10^0$
$10^1$
$10^2$
$10^3$
$10^4$
mmseqs2
minimap2
lexicmap
bbmap_skimmer
bowtie2
blastn
strobealign
mummer4
bowtie1

### C. 2 mismatches

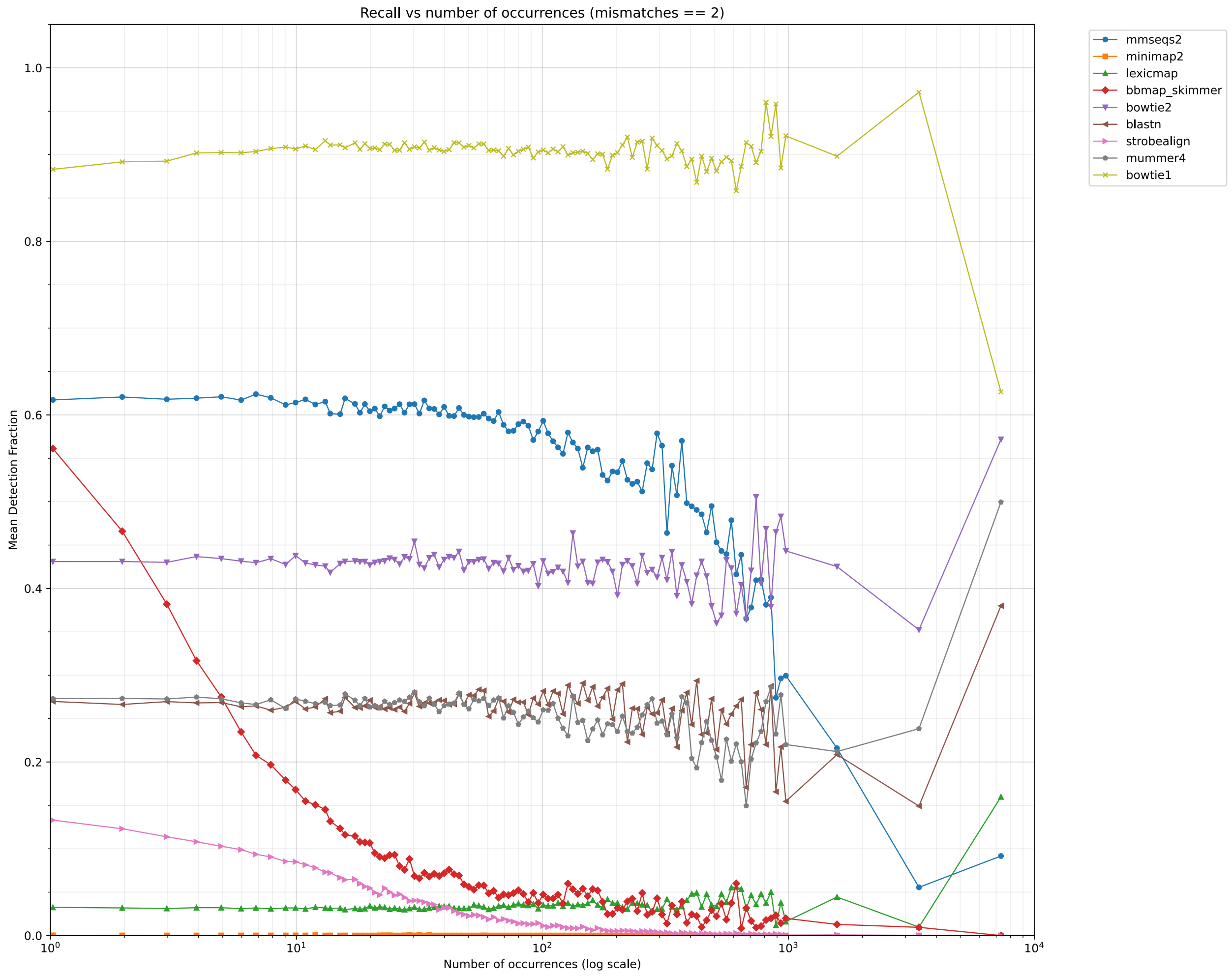

#### D. 3 mismatches

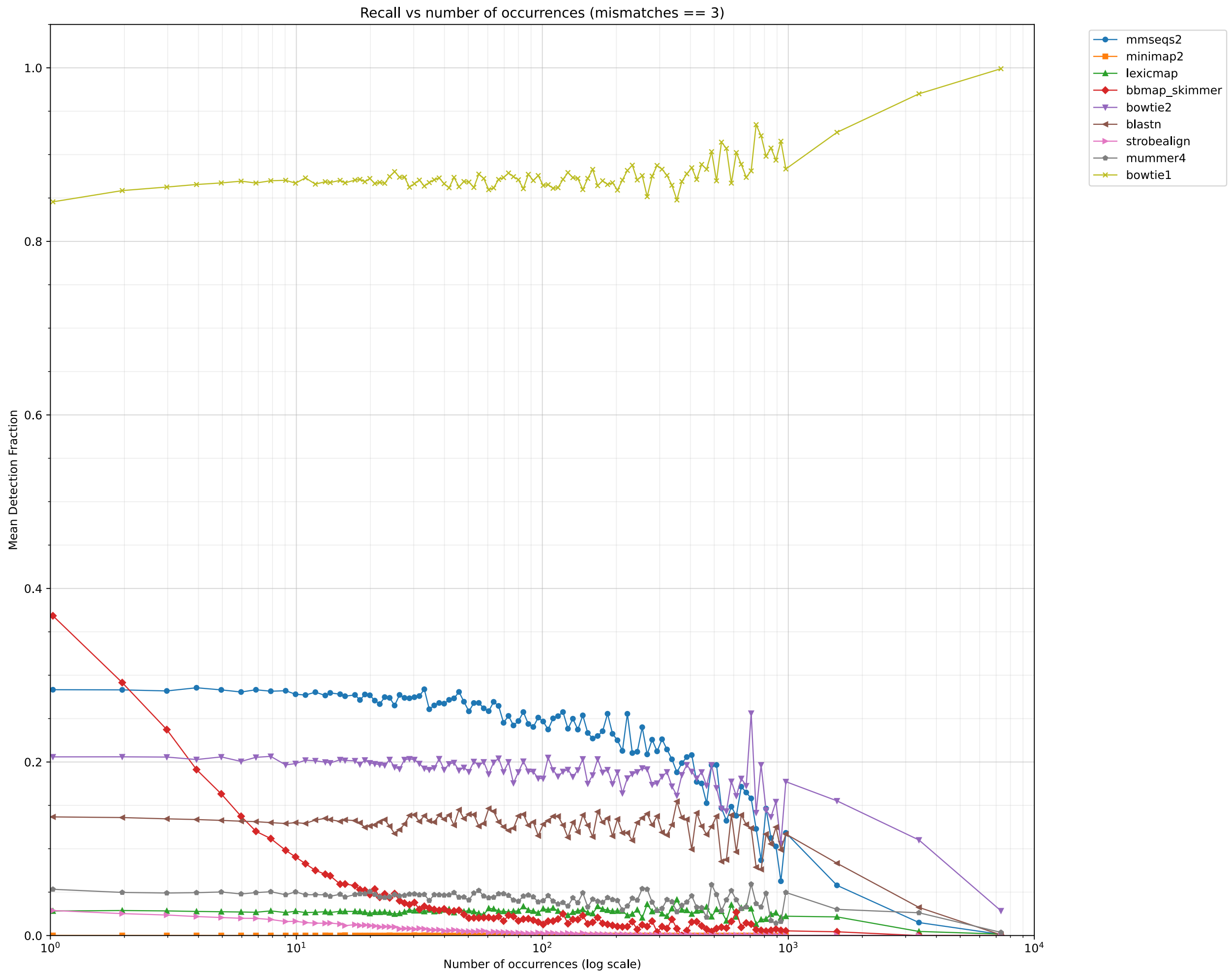


### Supplementary figure 6.

Distributions of spacers characteristics.
(A) Spacer size (length in bp)
(B) Mismatches observed in spacer alignments
(C) Spacer occurrence rate.

#### 1. IMG/VR4 dataset

**Note:** the horizontal axis is logarithmic.

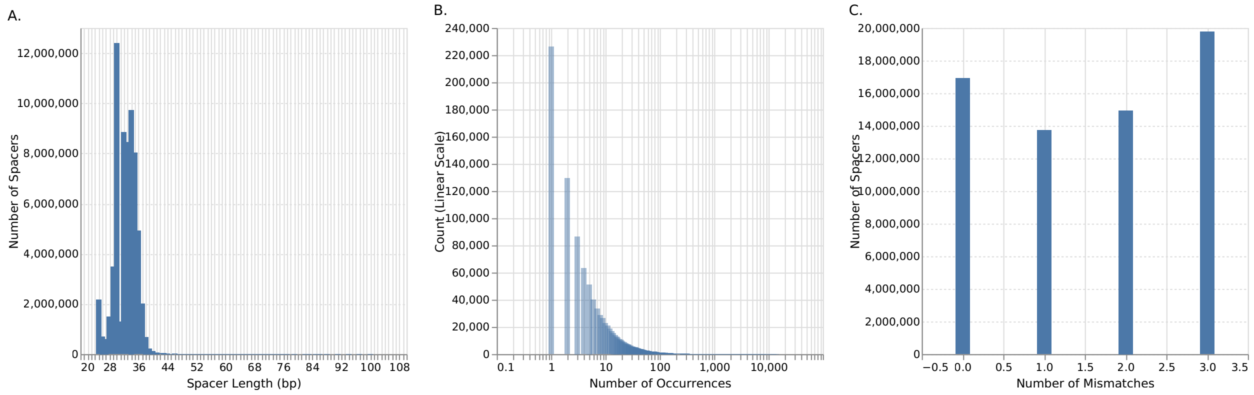


#### 2. Synthetic dataset

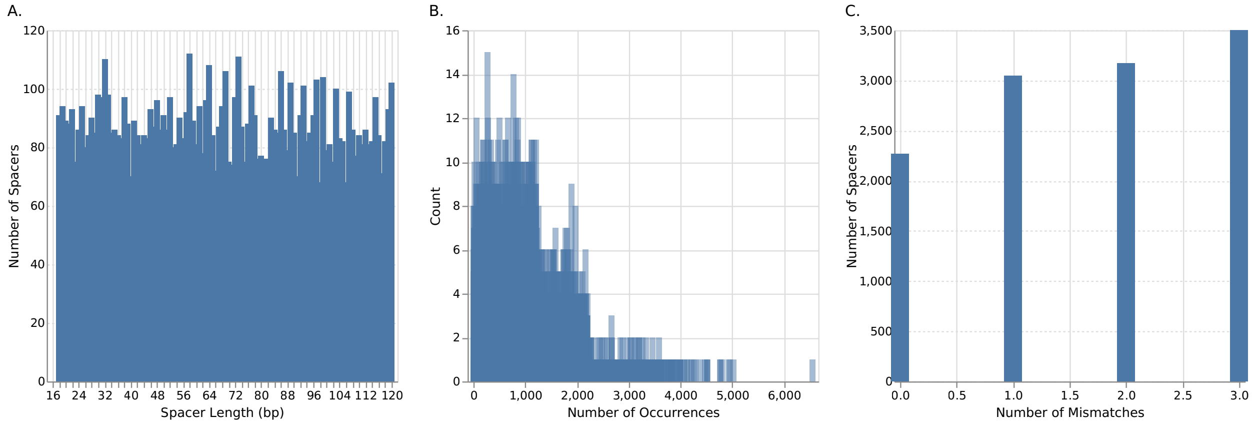


### Supplementary figure 7.

Upset plot of tool performance for IMG/VR4 dataset. The panels (from top to bottom) are sorted by number of mismatches. The sets in each panel are sorted from left to right by the set size. Each row in each panel represents a tool, and the vertical lines connecting the dots indicate the intersection of the connected tool results'.

#### A. 0 mismatches

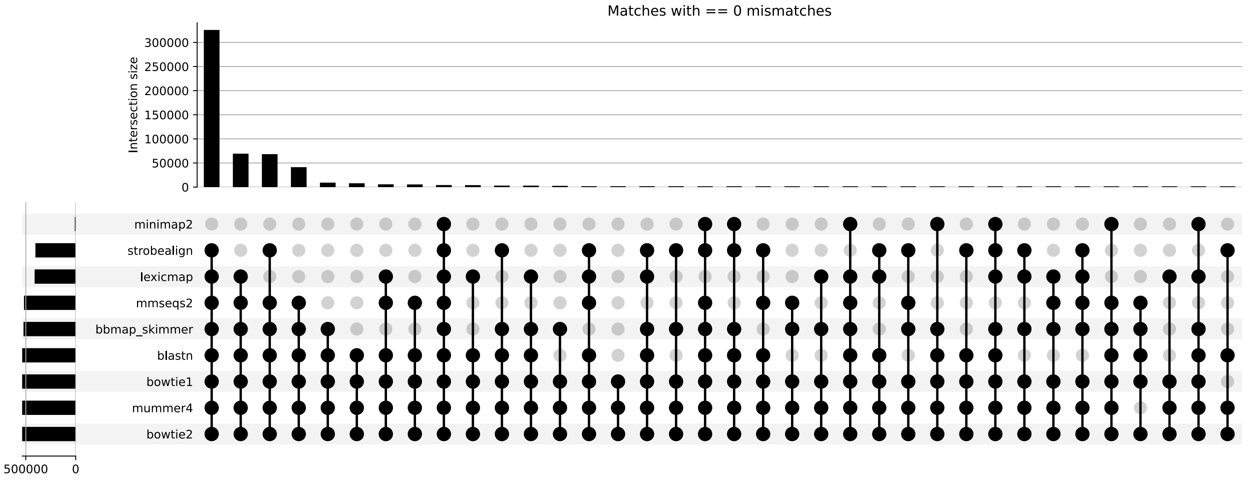

### B. 1 mismatch

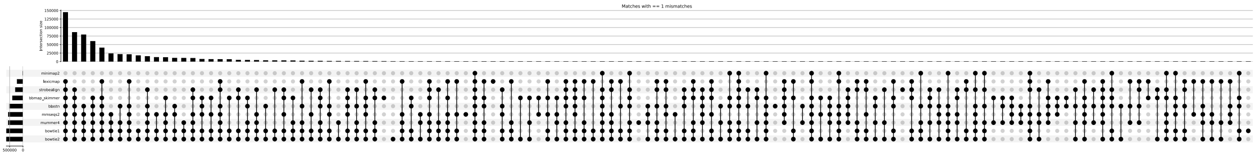


### C. 2 mismatches

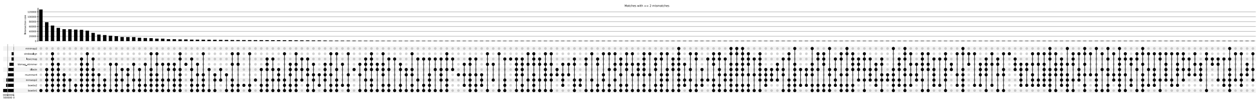

### D. 3 mismatches

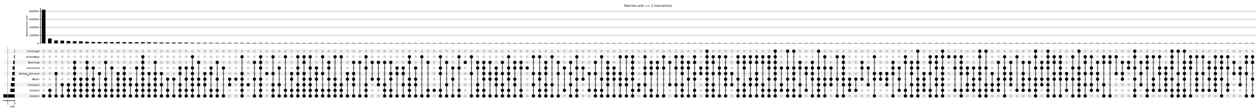
